# Postembryonic screen for mutations affecting spine development in zebrafish

**DOI:** 10.1101/2020.08.12.248716

**Authors:** Ryan S. Gray, Roberto Gonzalez, Sarah D. Ackerman, Ryoko Minowa, Johanna F. Griest, Melisa N. Bayrak, Benjamin Troutwine, Stephen Canter, Kelly R. Monk, Diane S. Sepich, Lilianna Solnica-Krezel

**Affiliations:** Department of Nutritional Sciences, Dell Pediatric Research Institute, University of Texas at Austin, Austin, Texas, USA; Department of Developmental Biology, Washington University School of Medicine, St. Louis, MO, USA; Vollum Institute, Oregon Health & Science University, 3181 SW Sam Jackson Park Rd., Portland, OR, 97239, USA.

## Abstract

The spinal vertebral column gives structural support for the adult body plan, protects the spinal cord, and provides muscle attachment and stability, which allows the animal to move within its environment. The development and maturation of the spine and its physiology involve the integration of multiple musculoskeletal tissues including bone, cartilage, and fibrocartilaginous joints, as well as innervation and control by the nervous system. One of the most common disorders of the spine in human is adolescent idiopathic scoliosis (AIS), which is characterized by the onset of an abnormal lateral curvature of the spine of <10° around adolescence, in otherwise healthy children. The genetic basis of AIS is largely unknown. Systematic genome-wide mutagenesis screens for embryonic phenotypes in zebrafish have been instrumental in the understanding of early patterning of embryonic tissues necessary to build and pattern the embryonic spine. However, the mechanisms required for postembryonic maturation and homeostasis of the spine remain poorly understood. Here we report the results from a small-scale forward genetic screen for adult-viable recessive and dominant mutant zebrafish, displaying overt morphological abnormalities of the adult spine. Germline mutations induced with *N*-ethyl *N*-nitrosourea (ENU) were transmitted and screened for dominant phenotypes in 1,229 F1 animals, and subsequently bred to homozygosity in F3 families, from these, 314 haploid genomes were screened for recessive phenotypes. We cumulatively found 39 adult-viable (3 dominant and 36 recessive) mutations each leading to a defect in the morphogenesis of the spine. The largest phenotypic group displayed larval onset axial curvatures, leading to whole-body scoliosis without vertebral dysplasia in adult fish. Pairwise complementation testing within this phenotypic group revealed at least 16 independent mutant loci. Using massively-parallel whole genome or whole exome sequencing and meiotic mapping we defined the molecular identity of several loci for larval onset whole-body scoliosis in zebrafish. We identified a new mutation in the *skolios*/*kinesin family member 6* (*kif6*) gene, causing neurodevelopmental and ependymal cilia defects in mouse and zebrafish. We also report several recessive alleles of the *scospondin* and *a disintegrin and metalloproteinase with thrombospondin motifs 9* (*adamts9*) genes, which all display defects in spine morphogenesis. Many of the alleles characterized thus far are non-synonymous mutations in known essential *scospondin* and *adamts9* genes. Our results provide evidence of monogenic traits that are critical for normal spine development in zebrafish, that may help to establish new candidate risk loci for spine disorders in humans.

## Introduction

Scoliosis is a medical condition characterized by complex three-dimensional deformity of the spine. The clinical classification of scoliosis is broadly defined by the time of onset of disease (e.g. congenital or after birth) and by whether the vertebrae are malformed. Scoliosis manifesting after birth is classified into two main groups: syndromic and idiopathic. Syndromic scoliosis is associated with known connective tissue disorders, such as Marfan or Ehlers-Danlos syndromes, and neurological conditions such as Rett syndrome or Horizontal Gaze Palsy, or as the consequence of acquired neuromuscular conditions, such as poliomyelitis. In contrast, idiopathic scoliosis (IS) occurs over a range of developmental ages in otherwise healthy children and without obvious malformation of the vertebrae. The most common form, called Adolescent Idiopathic Scoliosis (AIS), occurs typically during the growth spurt around adolescence (Wise et al., 2020).

The pathogenesis of AIS is hypothesized to be the result of many causes including biomechanical, neural/proprioceptive, and hormonal changes (Cheng et al., 2015). The genetic basis of AIS in human is only beginning to be understood. Whereas there is evidence of familial linkage with IS, and a high concordance in monozygotic twins (Wise et al., 2008); however, no monogenic traits have been demonstrated, even in familial cases (Miller, 2007). For this reason, it is generally accepted that IS likely represents a complex multifactorial inheritance disorder. The association of IS with several candidate loci have begun to implicate genes involved in connective tissue structure, bone formation/metabolism, melatonin signaling, onset of puberty, and growth; however, there are just as many, if not more, publications contradicting these reports for the same genes and pathways (Gorman et al., 2012).

Recently, large-scale genome-wide association studies (GWAS) of AIS patients have begun to implicate several independent loci in AIS (Ikegawa, 2016). For instance, variants in the *ADGRG6* (also called *GPR126*) locus are associated with AIS in a variety of ethnic backgrounds (Kou et al., 2013; Kou et al., 2018). Importantly, the conditional ablation of *Adgrg6* in osteochondral progenitor lineages of the spine generates a faithful model of AIS in mouse (Karner et al., 2015). The pathogenesis of scoliosis in this model is associated with changes in the intervertebral disc including decreased chondrogenic gene expression, increased pro-inflammatory signaling, and increased stiffness of the disc tissues prior to the onset of scoliosis (Liu et al., 2019a). Additional lines of evidence for the pathogenesis underlying scoliosis have been described as the result of reduced endochondral bone in the vertebral bodies of *Prmt5* conditional mutant mice, modeling aspects of infantile IS (Liu et al., 2019b); and after defects in the integration of proprioceptor input to the spinal cord in *Runx3* mutant mice, modeling AIS (Blecher et al., 2017). Altogether, these scoliosis mutant mouse models demonstrate that disruptions of structural elements and sensory-motor inputs of the spine may underlie the pathogenesis of IS in humans.

Given the present barriers to understanding the molecular genetics of IS in humans, we sought to generate adult-viable, genetically-tractable models of scoliosis using a forward genetic screen in zebrafish to identify conserved pathways, which regulate spine morphogenesis. Zebrafish have increasingly become an established model organism for the study of musculoskeletal development and associated disorders (Busse et al., 2019). Zebrafish have also become a premier animal model for mechanistic studies of scoliosis and vertebral malformations as a consequence of defects of the notochord sheath via analyses of mutations in the *collagen VIIIa1a* (*col8a1a*) gene (Gray et al., 2014) or defects in the biogenesis of notochord vacuoles (Bagwell et al., 2020; Ellis et al., 2013). Zebrafish also have been shown to model aspects of AIS in several independent genetic mutations, including in: *protein kinase 7* (*ptk7*) (Hayes et al., 2014); *kinesin family member 6* (*kif6*) (Buchan et al., 2014); and *signal transducer and activator of transcription 3* (*stat3*) (Liu et al., 2017). Studies of *stat3* and *ptk7* mutants uncovered a potential connection between inflammation and scoliosis (Liu et al., 2017; Van Gennip et al., 2018). Interestingly, in the case of the *ptk7* mutant zebrafish, scoliosis onset was delayed by treatment with anti-inflammation drugs, suggesting a potential therapeutic avenue for AIS in humans (Van Gennip et al., 2018).

Motile cilia in the brain and spinal cord and regulation of cerebrospinal fluid flow have been strongly implicated in the pathogenesis of AIS-like phenotypes in zebrafish. Mutations in zebrafish regulators of motile cilia development or physiology, such as *ptk7* (Grimes et al., 2016; Hayes et al., 2014), *kif6* (Buchan et al., 2014; Konjikusic et al., 2018), *cc2d2a* (Bachmann-Gagescu et al., 2011), *kurly/cfap298, ccdc40, ccdc151*, and *dyx1c1*(Grimes et al., 2016) result in scoliosis. Given these clear examples of adult-viable IS-like scoliosis in zebrafish, we reasoned that a forward genetic screen to identify additional recessive mutations causing scoliosis would greatly inform our understanding of the genetic causes of scoliosis in humans.

Here, we report the results of a small-scale forward genetic screen for ENU induced mutations in zebrafish, in which we identified, 39 adult mutants, comprising five phenotypic groups, which model aspects of IS or congenital scoliosis. Partial complementation testing of the phenotypic group comprising 16 mutations, which displays the onset of pathology and morphology of scoliosis reminiscent of human AIS, defined at least 10 loci. Whole genome or exome sequencing of these scoliosis mutants, identified new mutations in the *kif6* gene, which we previously implicated in zebrafish to model AIS (Buchan et al., 2014; Konjikusic et al., 2018), and two hypomorphic alleles of the *scospondin* gene, which encodes a critical protein component necessary for the development of the Reissner fiber (RF) and regulation of straightening of the body axis during embryonic development in zebrafish (Cantaut-Belarif et al., 2018; Troutwine et al., 2020). We also report several alleles of the *a disintegrin and metalloproteinase with thrombospondin motifs 9* (*adamts9*) gene, which is associated with a unique scoliosis morphology beginning in the caudal portion of the spine. Notably, the identified recessive scoliosis alleles are hypomorphic mutations of essential genes, underscoring the utility of a forward genetic approach for understanding the genetic susceptibility of spine disorders.

Our characterization of these new zebrafish mutant models of scoliosis support the established interdependency of the motile cilia function and RF assembly for spine morphogenesis. We also implicate a role of Adamts9 for scoliosis and demonstrate it is not due to obvious alterations in RF assembly. Altogether this screen underscores the heterogenous and multi-factorial nature of the genetic control of spine morphogenesis and these mutant zebrafish will allow us to distinguish distinct etiologies causing scoliosis and potentially inform new candidate risk loci of spine disorders in humans.

## Results

### Forward Genetic Screen for Mutations Affecting Spine Development

In zebrafish, large-scale systematic screens for recessive zygotic mutations have largely focused on embryonic and early-larval phenotypes carried out at stages when zebrafish do not require feeding (Driever et al., 1996). Only limited screens have been performed for mutations affecting postembryonic zebrafish development and were mostly focused on the identification of dominant mutations (Henke et al., 2017). The rationale for designing this screen was to facilitate the discovery of the molecular genetics of spine development and of adult-viable spine disorders and to provide tractable experimental models of these common human musculoskeletal disorders in zebrafish. To carry out a forward genetic screen for recessive mutations affecting spine development, we carried out ENU mutagenesis of F0 males as previously (Solnica-Krezel et al., 1994) with modifications described in Methods. The efficiency of mutagenesis was assessed by specific locus testing and revealed mutagenesis rates of 1/500 for the *albino*, 1/1000 for *golden* and 1/150 for *sparse* loci, about two times higher than in the Boston mutagenesis (*albino* 1/1225; *golden* 1/1170 and *sparse* 1/470) (Solnica-Krezel et al., 1994). We carried out a traditional breeding scheme, crossing F0 mutagenized males to wild-type (WT) females generating F1 animals, which were then outcrossed to WT fish to produce F2 families. Finally, crosses of F2 siblings produced F3 families, which were raised to 20-30 days post fertilization (dpf).

We screened 1,229 adult F1 animals to identify dominant mutations affecting spine morphology or body shape. All candidate dominant F1 mutations were backcrossed to an isogenic mapping strain (e.g. SJD or WIK) in order to confirm the dominant mutation in the F2 generation. We analyzed each of these crosses for reappearance of the original F1 phenotype at 30 dpf or later. All confirmed dominant mutants were processed for Alcian-Blue and Alizarin-Red skeletal staining in order to document the structure of the adult skeleton. Finally, all lines were cryopreserved (Matthews et al., 2018) for future analysis, as the main body of this study involved recessive mutations. We recovered three dominant mutations that caused shortened adult body length, incidence of vertebral fusions, without scoliosis (Group V, Table 1 and Supp. Fig.1), which were reminiscent of dominant zebrafish mutants recently reported (Henke et al., 2017). We did not discover any dominant mutations modeling AIS in this screen. Further characterization of these dominant mutations is not included within the scope of this report.

**Table 1.**
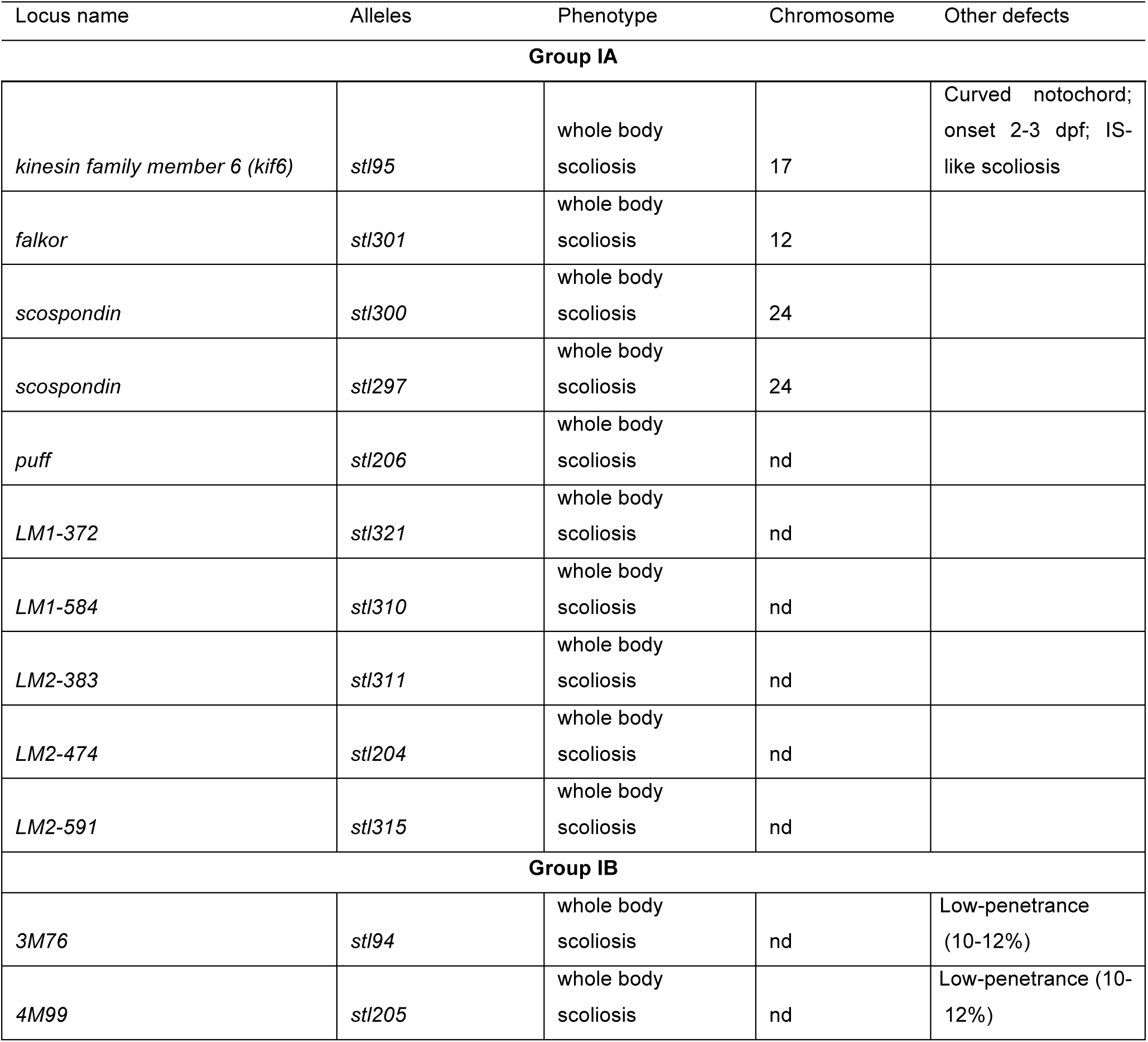
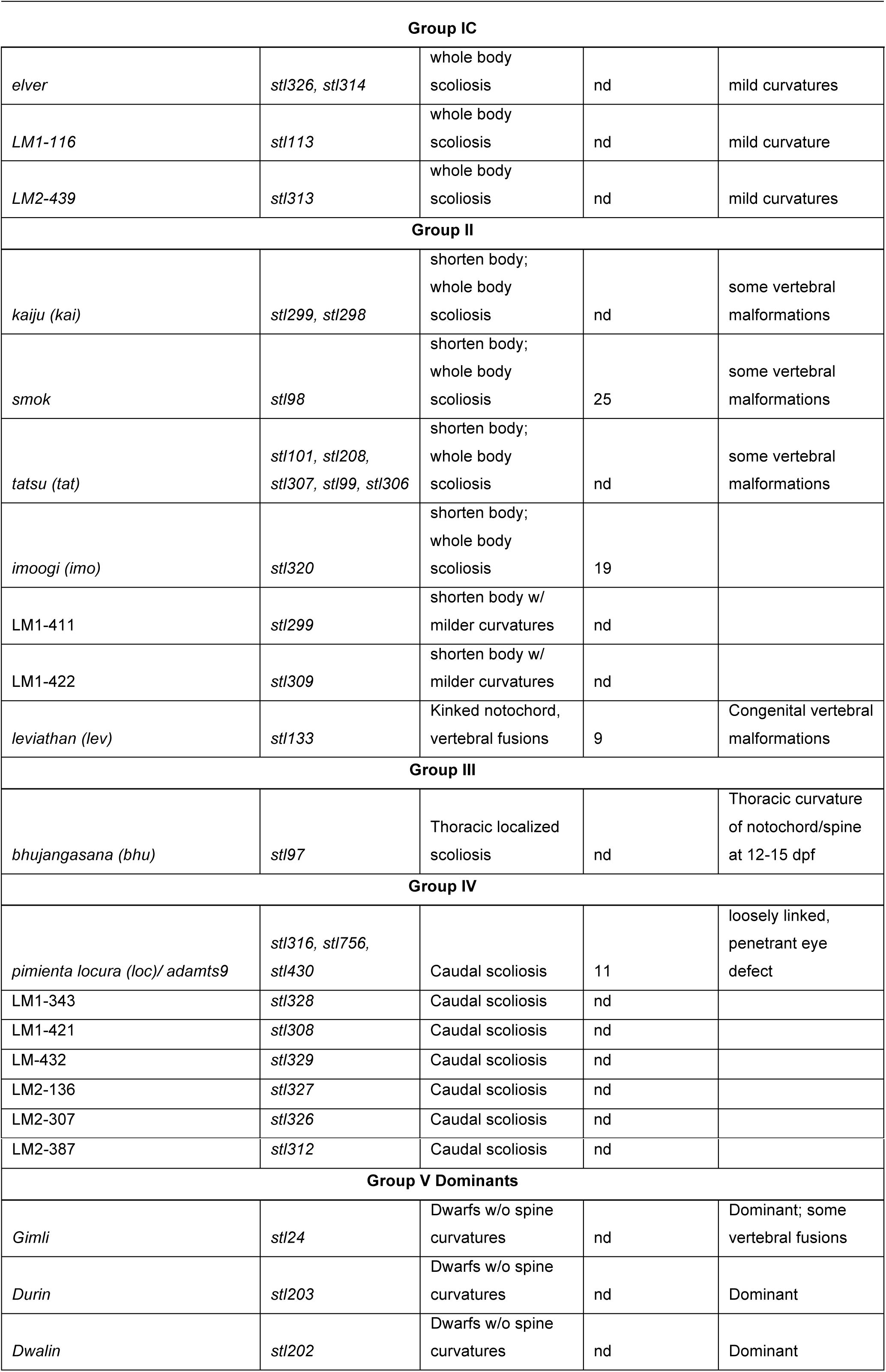
Zebrafish mutations affecting spine structure.

| Locus name | Alleles | Phenotype | Chromosome | Other defects |
| --- | --- | --- | --- | --- |
| **Group IA** |  |  |  |  |
| *kinesin family member 6 (kif6)* | *stl95* | whole body scoliosis | 17 | Curved notochord; onset 2-3 dpf; IS-like scoliosis |
| *falkor* | *stl301* | whole body scoliosis | 12 |  |
| *scospondin* | *stl300* | whole body scoliosis | 24 |  |
| *scospondin* | *stl297* | whole body scoliosis | 24 |  |
| *puff* | *stl206* | whole body scoliosis | nd |  |
| *LM1-372* | *stl321* | whole body scoliosis | nd |  |
| *LM1-584* | *stl310* | whole body scoliosis | nd |  |
| *LM2-383* | *stl311* | whole body scoliosis | nd |  |
| *LM2-474* | *stl204* | whole body scoliosis | nd |  |
| *LM2-591* | *stl315* | whole body scoliosis | nd |  |
| **Group IB** |  |  |  |  |
| *3M76* | *stl94* | whole body scoliosis | nd | Low-penetrance (10-12%) |
| *4M99* | *stl205* | whole body scoliosis | nd | Low-penetrance (10-12%) |

| Group IC |  |  |  |  |
| --- | --- | --- | --- | --- |
| *elver* | *stl326, stl314* | whole body scoliosis | nd | mild curvatures |
| *LM1-116* | *stl113* | whole body scoliosis | nd | mild curvature |
| *LM2-439* | *stl313* | whole body scoliosis | nd | mild curvatures |
| Group II |  |  |  |  |
| *kaiju (kai)* | *stl299, stl298* | shorten body;<br>whole body scoliosis | nd | some vertebral malformations |
| *smok* | *stl98* | shorten body;<br>whole body scoliosis | 25 | some vertebral malformations |
| *tatsu (tat)* | *stl101, stl208,*<br>*stl307, stl99, stl306* | shorten body;<br>whole body scoliosis | nd | some vertebral malformations |
| *imoogi (imo)* | *stl320* | shorten body;<br>whole body scoliosis | 19 |  |
| *LM1-411* | *stl299* | shorten body w/<br>milder curvatures | nd |  |
| *LM1-422* | *stl309* | shorten body w/<br>milder curvatures | nd |  |
| *leviathan (lev)* | *stl133* | Kinked notochord,<br>vertebral fusions | 9 | Congenital vertebral malformations |
| Group III |  |  |  |  |
| *bhujangasana (bhu)* | *stl97* | Thoracic localized scoliosis | nd | Thoracic curvature of notochord/spine at 12-15 dpf |
| Group IV |  |  |  |  |
| *pimienta locura (loc)/ adamts9* | *stl316, stl756,*<br>*stl430* | Caudal scoliosis | 11 | loosely linked, penetrant eye defect |
| *LM1-343* | *stl328* | Caudal scoliosis | nd |  |
| *LM1-421* | *stl308* | Caudal scoliosis | nd |  |
| *LM-432* | *stl329* | Caudal scoliosis | nd |  |
| *LM2-136* | *stl327* | Caudal scoliosis | nd |  |
| *LM2-307* | *stl326* | Caudal scoliosis | nd |  |
| LM2-387 | *stl312* | Caudal scoliosis | nd |  |
| **Group V Dominants** |  |  |  |  |
| *Gimli* | *stl24* | Dwarfs w/o spine curvatures | nd | Dominant; some vertebral fusions |
| *Durin* | *stl203* | Dwarfs w/o spine curvatures | nd | Dominant |
| *Dwalin* | *stl202* | Dwarfs w/o spine curvatures | nd | Dominant |

With regards to recessive mutations, 671 F3 families were screened between 20-30 dpf (ranging from 6.4-6.9 mm on average) for obvious spine/body plan defects, yet allowing enough fitness to survive larval development within a population of phenotypically WT siblings. Our cutoff for calling a potential mutant expected to manifest at 25% at full penetrance, was at ≥10 % penetrance of the total clutch. Such phenotypic putative recessive mutants were subsequently isolated for rearing to adulthood, to increase the likelihood of survival to sexual maturity for backcrossing to the SAT background. This strategy allowed us to isolate and expand the potential recessive mutation and further facilitate the verification of their recessive nature in the F5 generation. Genomic DNA was collected from some mutants and their WT siblings from a single F5 family for future genomic analysis. Concurrently, all recessive mutations were also analyzed at 5 dpf for embryonic phenotypes, in particular to identify abnormalities of notochord development, axial curvatures, and atypical swimming behaviors. At 30 dpf, we processed representative WT and phenotypic individuals for skeletal preparations using Alcian-Blue and Alizarin-Red to label cartilage and mineralized matrix.

### Phenotypic classes of recessive mutations

We based phenotypic classification of the 36 identified recessive mutations on the distribution and type of scoliotic bends, and overall body length. Using these metrics, we distinguished four distinct phenotypic groups: Group I: whole-body scoliosis and largely normal length, without vertebral malformations (Fig. 1B and 2B); Group II: shorten body, with and without vertebral malformations (Fig. 1C and 2C); Group III: thoracic-localized lordosis (Fig. 1D and 2D); and Group IV: caudal-localized scoliosis (Fig. 1E and 2E); (Table1).

**Figure 1:**
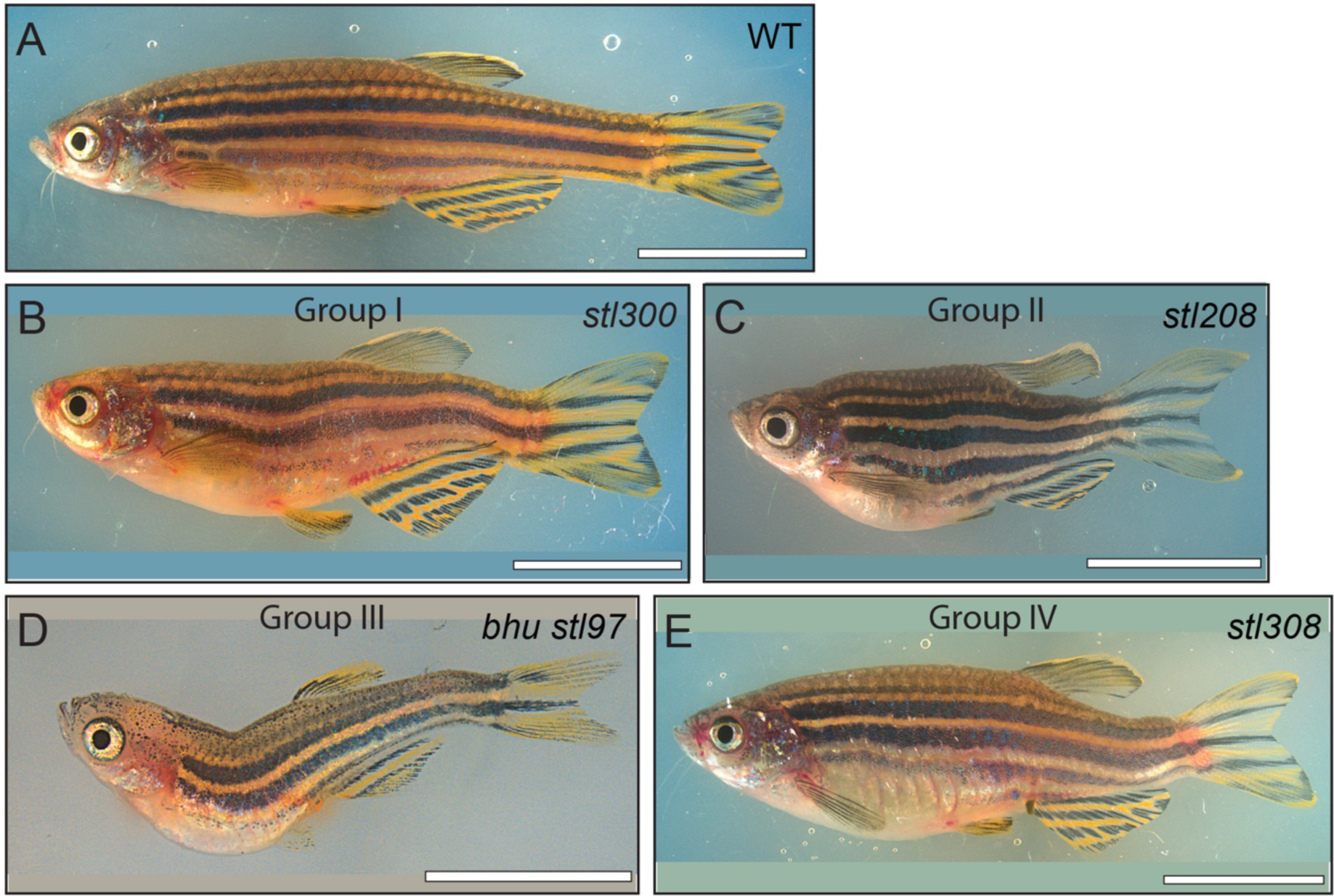
Representative phenotypes of the spine defect mutant classes at post-metamorphic stages. Bright field pictures of wild type **(A)** and representative mutants (B-E), displaying whole-body scoliosis **(B**, Group I), dwarf or shorten body plan **(C**, Group II), thoracic-localized scoliosis **(D**, Group III), or caudal-localized scoliosis **(E**, Group IV). Scale bars, 1 cm, except D, which is 5 mm.

**Figure 2:**
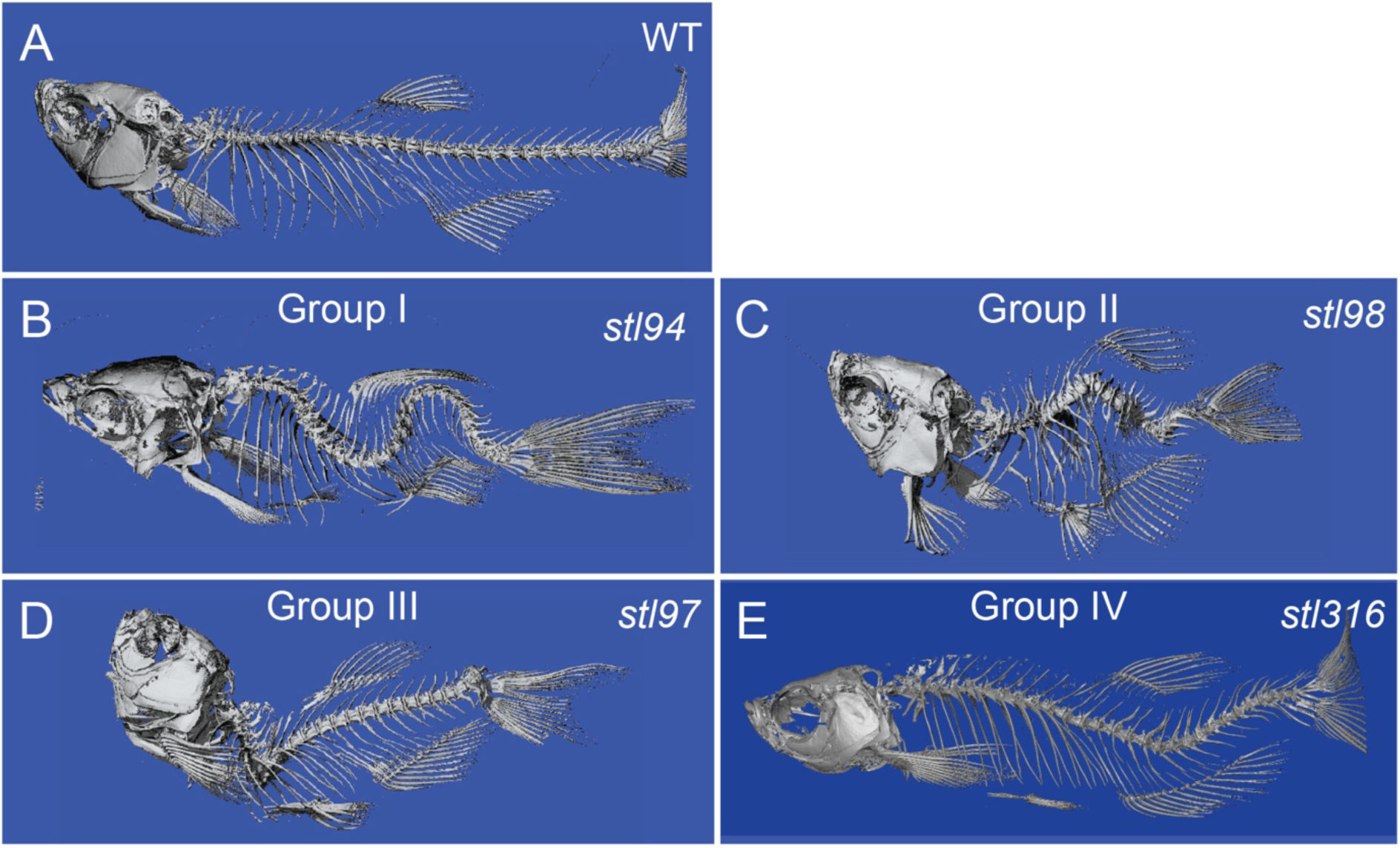
Micro-computed tomography imaging of the representative spine defect mutant classes. µCT imaging of wild type **(A)** and representative mutants **(B-E)**, displaying whole-body scoliosis (*stl94*) **(B**, Group I), dwarf or shorten body plan and vertebral body defects (*stl98*) **(C**, Group II), thoracic-localized scoliosis (*bhu*^*stl97*^) **(D**, Group III), and caudal-localized scoliosis *loc*^*stl316*^ **(E**, Group IV).

To begin complementation testing, we selected 16 independent lines from Group IA, IC, and Group II for comprehensive pair-wise complementation testing, using non-phenotypic heterozygous carriers of each mutation. After each pairwise cross, we assayed the progeny for obvious body curvatures at 15 and 30 dpf (Fig. 1). From these crosses, we defined six complementation groups: one comprising two mutations in Group IA (*stl297* and *stl300*); one comprising two mutations in Group 1C (*stl314* and *stl326*); one showing interaction between Group IA and II mutations (*stl321* and *stl98*); and three groups displaying complex interactions between several Group II mutations (*stl298, stl101, stl208, stl307, stl306*, and *stl99*). In contrast, several Group I mutations tested did complement one another, suggesting that these phenotypes involve multiple independent loci. Due to the logistics of carrying out complementation testing out to 30 dpf, we were not able to perform comprehensive complementation testing for all mutations identified in this screen. Ongoing mapping efforts provide a parallel approach to test allelism of these mutations (Table 1).

### Group I: mutants displaying whole-body scoliosis, without vertebral malformations

The largest phenotypic group we discovered was composed of 16 recessive mutations, and may possess the most potential for modeling IS in humans (Fig 1B and 2B). This group was further divided into three subgroups based on the penetrance and expressivity of spine defect. Mutations in Group IA were fully penetrant, being manifested as spine curvatures along the entire body length by 100% of homozygous mutants, Group IB mutations were incompletely penetrant, but also generating spine curvatures along the entire body, and Group IC mutations were fully penetrant but generated milder curves.

### Group II: mutants displaying a shorten body axis with scoliosis

This group was the second largest phenotypic group uncovered in our screen, with 12 mutations (Fig 1C and 2C). These mutations cause scoliosis and mostly shorten the body length compared to Group I mutants or their WT siblings. In some cases, we observed vertebral malformations, however at present we have not fully characterized the timing of these defects or whether they are primary or secondary to scoliosis. Based on the complex results from the complementation testing of these mutants, further mapping and molecular characterization studies will be needed to determine whether these mutations affect one or more loci.

We identified a single group II mutation *stl133* associated with severe embryonic defects (e.g. bending and kinking) of the notochord and shorten body length and vertebral body defects in adult fish. This mutation displayed an identical phenotype as mutations in the *leviathan/collagen8a1a* (*col8a1a*) locus (Gray et al., 2014). Complementation testing between carriers of *stl133* mutation and a known *col8a1a*^*vu41*^ mutation, produced mutant progeny at an expected Mendelian frequency, which supports the notion that this mutation represents an uncharacterized mutation in *col8a1a*. This underscores the importance of *col8a1a* in the development of the notochord for adult spine morphogenesis.

### Group III: mutants displaying thoracic localized spine curvature

This group consists of one recessive mutation *bhujangasana/bhu*^*stl97/stl97*^, manifested as a larval-onset lordosis of the thoracic spine, which was apparent around 15 dpf (∼SL6) (Fig. 1D, 2D, 3B-B’’). This initial defect of larval *bhu*^*stl97/stl97*^ mutant spine sometimes progressed towards a severe lordosis, with rotation of the defective vertebral units in adult mutants (Fig. 2D).

**Figure 3:**
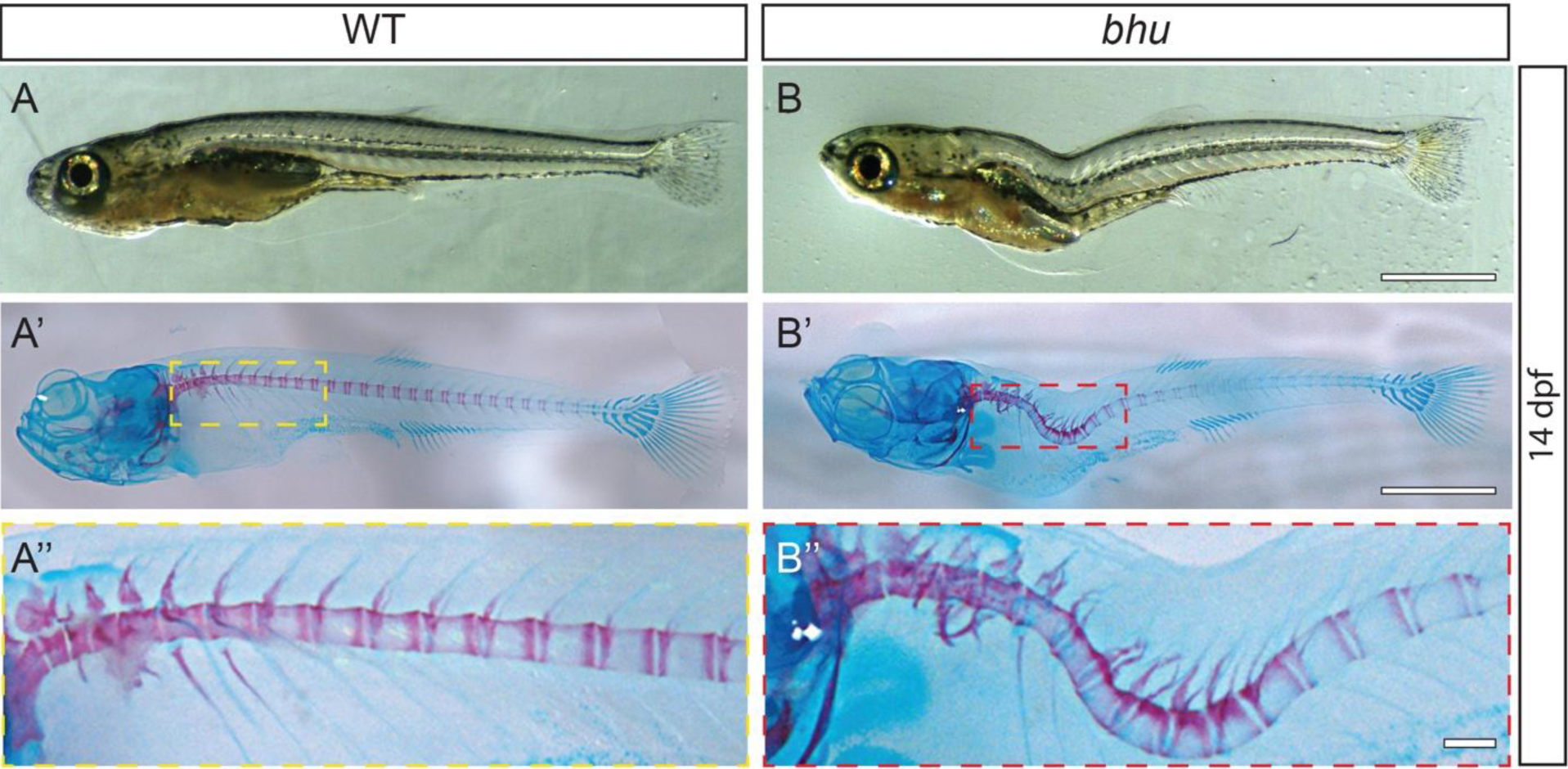
The bhu^stl97^ mutation generates a thoracic-localized lordosis during larval development. Bright field imaging of representative WT **(A-A”)** and *bhu*^*stl97/stl97*^ **(B-B”)** at 14 dpf shown as both unstained **(A, B)** and Alizarin-Red and Alcian-Blue stained larvae **(A’-A’’, B’-B’’)**. Scale bars, 1 mm.

### Group IV: mutants displaying caudal localized spine curvature

This group consists of seven mutations (Table 1, Fig. 1E, 2E), resulting in the formation of a single dorsalward bend of the caudal portion of the spine. We did not undertake complementation testing with this group. The genomic sequencing and characterization of one of members of this group, *stl316*, is detailed below.

### Group IA: *stl95* is a novel missense allele of the *skolios/kif6 gene*

To determine the molecular basis of a few of these mutant phenotypes we used two complementary approaches, standard SSLP bulked-segregant mapping (Knapik et al., 1998), whole-genome sequencing (WGS) and more recently, a zebrafish specific exome capture sequencing strategy (see Methods), to map mutant loci as revealed by regions of homozygosity and the included candidate mutations (Henke et al., 2013a; Henke et al., 2013b; Sanchez et al., 2017). Homozygous *stl95* mutants display mild to severe curvatures of the notochord at 5 dpf (Fig. 4A) followed by whole-body scoliosis without vertebral malformations in larval and adult fish (Fig. 4B, C). Evidence from bulk segregant mapping and WGS mapping identified a candidate region on Chromosome 17 (Fig. 4D). Previous work from our group uncovered both ENU-induced and Transcription activator-like effector nuclease (TALEN)-derived nonsense mutations of the *kif6* gene, also residing on Chromosome 17, and which display similar phenotypes as those observed in *stl95* mutants (Buchan et al., 2014; Konjikusic et al., 2018). To test whether the *stl95* mutation represented a new *kif6* allele we performed complementation testing with a previously reported TALEN-derived *kif6*^*Δ8*^ insertion-deletion (INDEL) mutation (Buchan et al., 2014). Multiple crosses of a heterozygous *stl95* mutant to a homozygous *kif6*^*Δ8/Δ8*^ mutant revealed that the two mutations failed to complement Group IA scoliosis phenotypes, demonstrating that *stl95* is a new allele of *kif6* (Table 2). Accordingly, analysis of WGS data revealed a homozygous T728A transversion mutation (ENSDART00000103662.6) in 100% of the reads derived from the *kif6*^*stl95*^ mutant pool. We observed the T728A mutation in 38% of the WGS reads from gDNA isolated from the aphenotypic sibling pool, consistent with the predicted 33% representation of the mutation in this group. This mutation is predicted to result in a L243Q amino acid change located in the putative switch II domain of the Kif6 motor domain (Marx et al., 2009). Clustal W alignment (Jeanmougin et al., 1998) of KIF6 proteins from multiple species demonstrates that this Leucine residue is highly conserved.

**Table 2:**
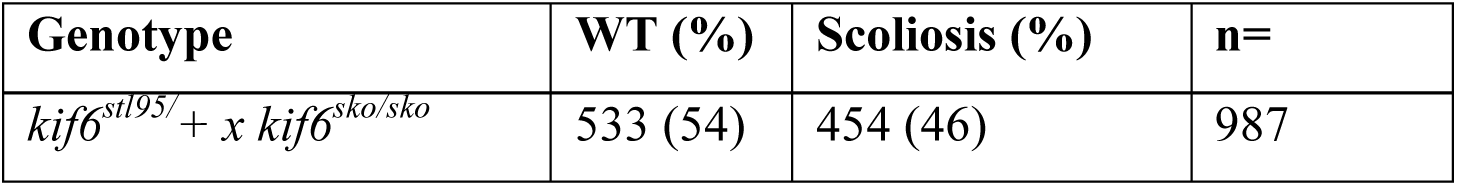
*kif6* complementation crosses.

**Figure 4:**
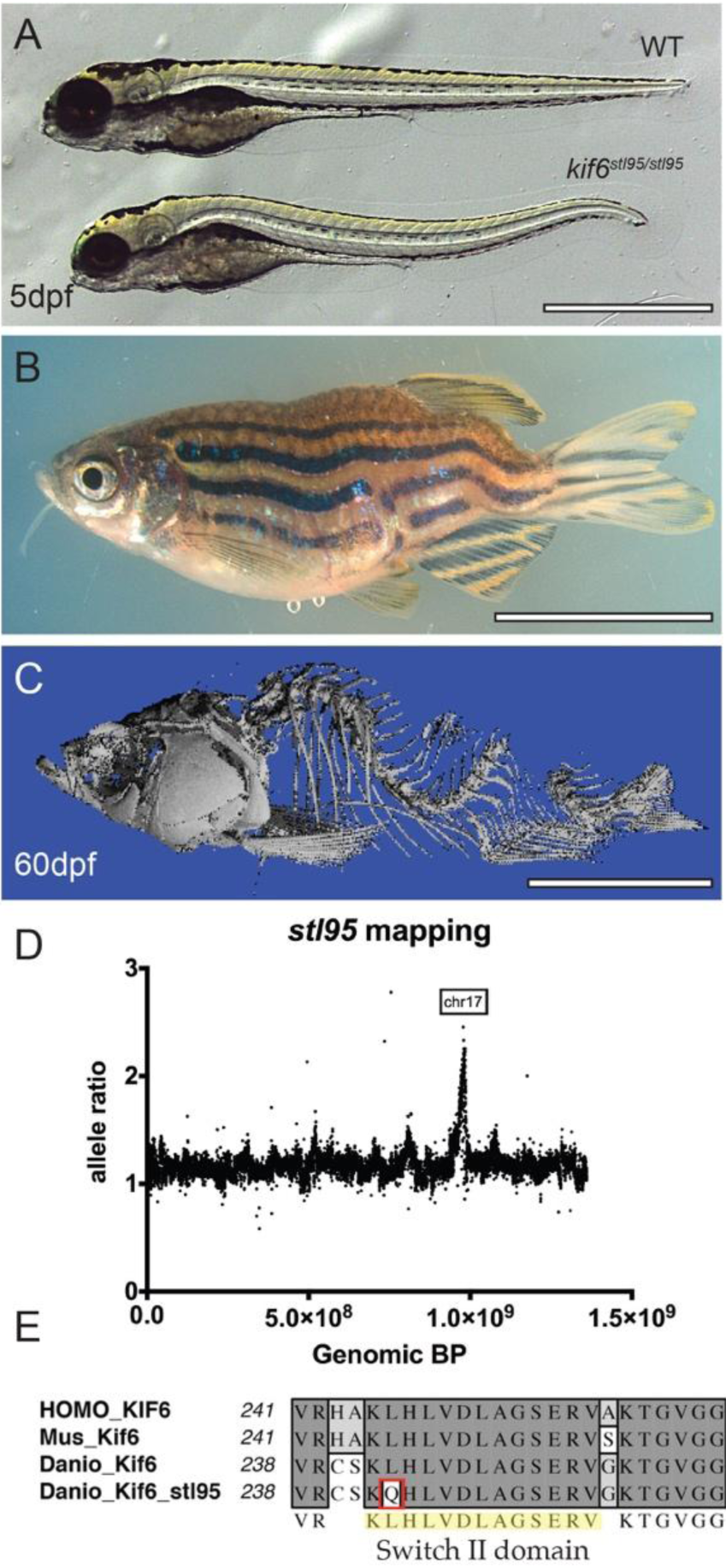
The *kif6*^*stl95*^ mutant displays early notochord bending and larval onset scoliosis with rotation, without vertebral malformations. Bright field of representative sibling WT and mutant *kif6*^*stl95/stl95*^ phenotypes at 5 dpf **(A)** and mutant adult at 60 dpf **(B)**. μCT imaging of a *kif6*^*stl95/stl95*^ mutant at 60 dpf **(C)**. Region of homozygosity-based mapping graphed as mutant allele ratio over the total genomic distance shows a peak of homozygosity at Chromosome 17 **(D)**. Multi-species alignment of the switch II domain of Kif6 proteins illustrating the amino acid change in the *kif6*^*stl95/stl95*^ mutant to Glutamine from a well-conserved Leucine residue **(E)**. Scale bars (A), 1 mm and (B, C), 1 cm.

Given that the T728A mutation in *kif6*^*stl95/stl95*^ alters a conserved motor domain residue and caused scoliosis in zebrafish, (Fig. 3E) we wanted to test if additional human mutations, in conserved kinesin residues, were also associated with disease. We requested *KIF6* variants from the Undiagnosed Disease Network (Baylor Miraca Genetics Laboratories) and identified a single c.886C>A/ p.P296T mutation in combination with a potentially damaging splice site mutation (c.1181+1G>A), associated with dopa responsive dystonia, epilepsy, and anxiety in a single patient. We found that this proline residue was conserved within the motor domain of the *Danio rerio* Kif6 protein (p.P293) (Supp. Fig. 2A), and within the motor domains of several other kinesin family proteins (not shown). To test the pathogenicity of the proline variant we employed a genome editing strategy using CRISPR/Cas9 to introduce this p.P293T mutation into the zebrafish *kif6* gene using an oligonucleotide to template the desired mutation, which also introduced a fortuitous EcoRI endonuclease site (Supp. Fig. 2A). By screening sperm samples by PCR amplification with the edit-specific PCR primers, we identified a single founder male. Screening its offspring by EcoRI digest identified a carrier heterozygous for the p.P293T allele (hereafter called *kif6*^*dp20*^), which was confirmed by Sanger sequencing (Supp. Fig. 2C, D). Homozygous *kif6*^*dp20/dp20*^ mutant zebrafish displayed scoliosis in adult mutants (Supp. Fig. 2E, F), which was phenotypically identical to that observed in *kif6*^*stl95/stl95*^ and *kif6*^*Δ8/Δ8*^ adult mutants (Buchan et al., 2014). These results underscore the critical role of *kif6* in spine morphogenesis in zebrafish and demonstrate that non-synonymous mutations of evolutionarily conserved amino acids in Kinesin proteins, in particular within the motor domain, should be promoted for further analysis of undiagnosed human disease, which can manifest in diverse pathologies (Chiba et al., 2019).

### Additional Group I mutant loci map to Chromosomes 12 and 24

Using bulked segregant mapping we were able to identify additional loci associated with larval onset scoliosis phenotype in zebrafish including (i) *falkor*^*stl301*^(Supp. Fig. 3A, B) (z4188, fa99f08, z8464) on linkage group (LG)12 (Supp. Fig. 3C); and (ii) two mutations, which failed to complement one another, *stl297* and *stl300* (z249, z9325, z6188), on LG24. Subsequently, we mapped the *falkor*^*stl301*^ mutation using our WGS pipeline to Chromosome 12 (Supp Fig. 3D), however the causative mutation remains undefined. Using WGS and WES we determined that both *stl297* and *stl300* were associated with non-synonymous mutations in the *scospondin* gene, which are described in greater detail in another manuscript (Troutwine et al., 2020). The remaining mutations have been cryopreserved and are being systematically introduced into our exome sequencing pipeline to facilitate mapping and identification of the causative mutations.

### The Group IV *pimienta locura/loc*^*stl316/stl316*^ mutation affects *adamts9*

We found several mutants, which displayed spine curvature that started in the caudal portion of the body, hereafter classified as caudal scoliosis. One of the members of this group, *pimienta locura/loc*^*stl316/stl316*^, exhibited larval onset lordosis around 15-18 dpf (∼SL6) (Fig. 5 A). This resulted in an overt caudal-localized body curvature phenotype in adult fish (Fig. 5 B, C) and scoliosis of the underlying spine (Fig. 5 D, E). This mutant also manifested an eye defect with low penetrance and variable expressivity, including dilated pupils and smaller misshapen eyes (Supp. Fig. 4). Barbels, which are whisker-like sensory organs near the mouth (red arrow, Fig. 5 D’) (LeClair and Topczewski, 2010), were consistently kinky or curly in *loc*^*stl316/stl316*^ mutant adults in comparison to non-phenotypic siblings (red arrow, Fig. 5E’).

**Figure 5:**
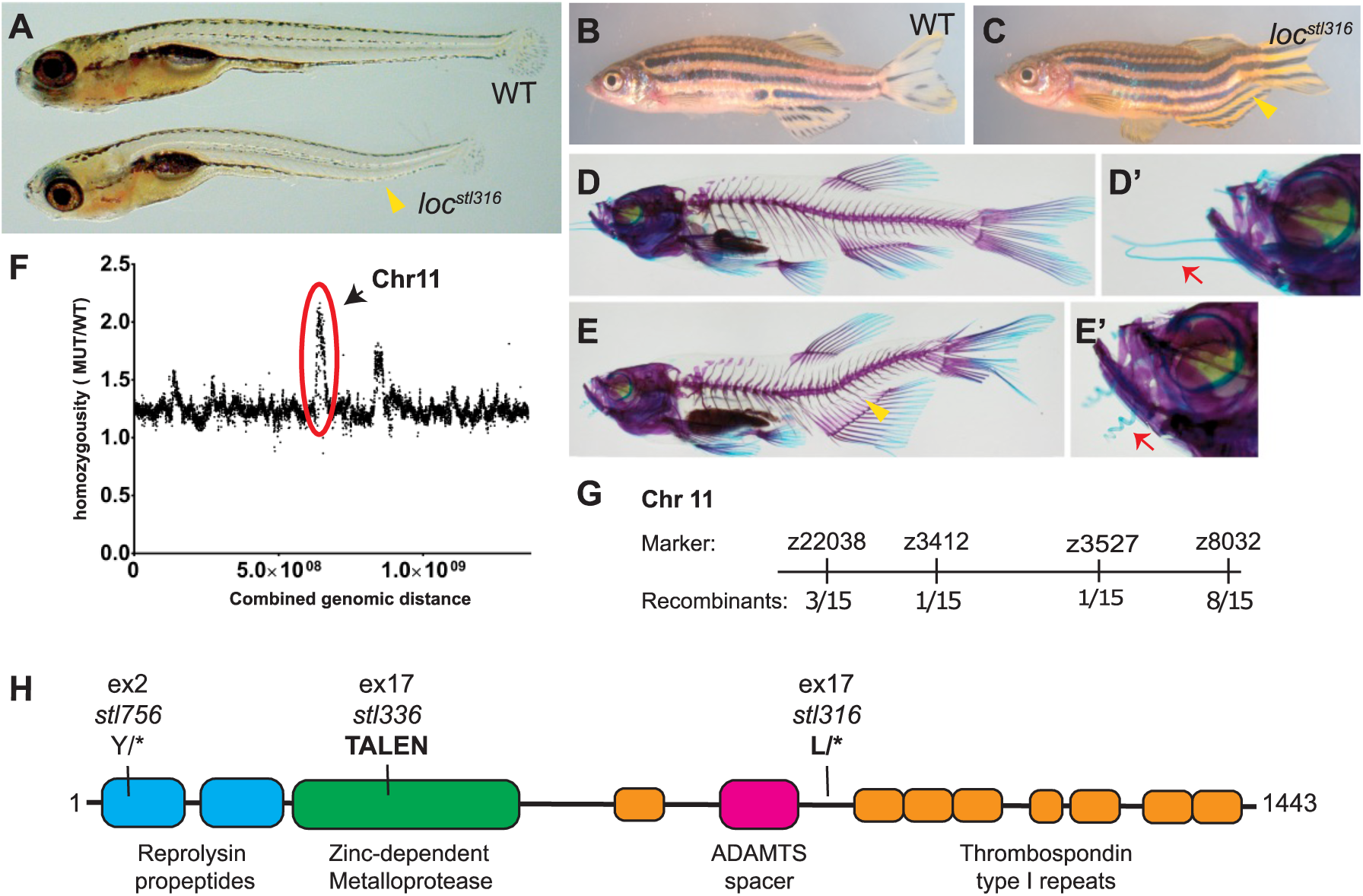
The *pimienta locura/loc*^*stl97*^ mutation is predicted to generate a truncated Adamts9 protein. Bright field of representative sibling WT and mutant *loc*^*stl316/stl316*^ phenotypes at 5 dpf **(A)** and 60 dpf **(B, C)** and Alizarin-Red and Alcian-Blue skeletal preparations of WT **(D, D’)** and *loc*^*stl316/stl316*^ mutants **(E, E’)** highlighting the caudal-localized curvature of the notochord and spine (yellow arrowhead). Close-ups of zebrafish faces highlighting phenotypic differences in barbel morphology (red arrows) normally straight in the wild type **(D’)** and curly or kinky in *loc*^*stl316/stl316*^ mutants **(E’)**. Homozygosity based mapping graphed as mutant allele ratio over the total genomic distance shows a major peak of homozygosity at Chromosome 11 and a minor peak at Chromosome 15 **(F)**. The *loc*^*stl316*^ lesion was mapped to a region of Chromosome 11 between markers z3412 and z3527 (number of recombinants are denoted at each marker) representing a large ∼26Mb region annotated for 519 genes in the Zv10 build. A schematic of the predicted 1443-amino acid protein, showing the locations of the three novel *adamts9* mutant alleles in this study **(H)**.

In order to determine the molecular basis of the *pimienta locura/loc*^*stl316/stl316*^ phenotypes, we utilized WGS mapping and identified two region-of-homozygosity peaks on Chromosome 11 and Chromosome 15 (Fig. 5 F). Using SSLP based meiotic mapping (Knapik et al., 1998), we determined the interval most linked to the *loc*^*stl316*^ located on LG 11 (Fig. 5 G). Within this interval, we identified a candidate nonsense mutation in the *adamts9* gene (ENSDART00000109440.4), encoded by a T2639A transversion mutation, which was present in 100% of the reads derived from the mutant pool, but was underrepresented (47%) in the non-segregating sibling pool. This nonsense mutation is predicted to result in an L880STOP C-terminal of the putative ADAMTS9 spacer domain and truncating the predicted C-terminal Thrombospondin type I repeats (Fig. 5 H) (Kelwick et al., 2015).

In order to further test that the *loc*^*stl316/stl316*^ (subsequently called *adamts9*^*stl316*^) mutant phenotype is caused by the L880STOP mutation in *adamts9* we performed complementation testing using another candidate nonsense mutation in *adamts9* (*stl756*), which we uncovered in our concurrent forward-genetic screen as an embryonic lethal mutation (Fig. 5H). The *stl756* mutation was also mapped by WGS to Chromosome 11 and a homozygous C333G mutation located in the *adamts9* gene was identified in the mutant pools, while being underrepresented in the sibling pools. The *stl756* mutation is predicted to generate a Y111STOP early in the N-terminal reprolysin propeptide domain of Adamts9 (Fig. 5H). The *adamts9*^*stl756*^ allele introduces a MseI restriction endonuclease site allowing for robust restriction polymorphism genotyping (Supp. Fig. 5A). In a parallel approach, we used TALEN genome editing to generate a novel INDEL mutation predicted to disrupt the 5’ end of the *adamts9* gene (Fig. 5H). The resulting, *adamts9*^*stl336*^ allele, is a complex INDEL mutation, which eliminates a conserved *HphI* restriction site in the *adamts9* gene (Supp. Fig. 5B). In contrast to robust adult-viable scoliosis observed in *loc*^*stl316*^ mutant zebrafish, both *adamts9*^*stl756/stl756*^ and *adamts9*^*stl336/stl336*^ mutants mostly exhibited embryonic lethality (∼5 dpf) with pleiotropic defects, which precluded analysis of the maturing spine.

In order to better characterize the spine defects associated with *adamts9* mutations we followed progeny derived from heterozygous *adamts9*^*stl756/+*^ parents to define the onset of developmental defects. At 6 dpf, *adamts9*^*stl756/stl756*^ mutants showed normal morphology, however most displayed poorly inflated swim bladders (n=20/20 mutants) (Supp. Fig. 6B). At 20 dpf, both *adamts9*^*stl756/stl756*^ and *adamts9*^*stl336/stl336*^ mutants (n=10/10 mutants) were much smaller than siblings, displayed more severe body curvatures (Supp. Fig. 6C, D) than was observed for either *adamts9*^*stl316/stl316*^ or maternal-zygotic (MZ) *adamts9*^*stl316/stl316*^ mutants (Table 3). We next genotyped 40 fish per developmental time point from several independent clutches derived from heterozygous *adamts9*^*stl756/+*^ parents. As development progressed, we observed a marked decrease in the number of surviving *adamts9*^*stl756/stl756*^ mutants in independent families beginning at 6 dpf. By 29 dpf only 3% of animals genotyped as mutants (n=40 animals/time point; *p<*.*0001*, Fisher’s exact test). The decrease in survivorship was correlated with the timing of putting larval fish on flowing water. We hypothesize the mutants were unable to swim and grow in a turbulent environment. To increase survivorship of *adamts9*^*stl756/stl756*^ mutants, we raised several families without flowing water through adulthood. At 8.5 months post fertilization, *adamts9*^*stl756/stl756*^ mutants (n=6) displayed agape mouth, small eyes and caudal spinal curvatures, as well reduced size compared to WT siblings (n=8) (Supp. Fig. 6 E, F). Similar observations were made for *adamts9*^*stl336/stl336*^ mutants (data not shown).

**Table 3:** *adamts9* complementation crosses.

| Cross | n (%) wild type | n (%) mild | n (%) severe | n= |
| --- | --- | --- | --- | --- |
| *stl316/+ x stl316/+* | 262 (84.2) | 36 (11.6) | 13 (4.2) | 311 |
| *MZstl316/+ x MZstl316/+* | 262 (74.6) | 83 (23.6) | 6 (1.7) | 351 |
| *stl316/+ x stl756/+* | 259 (81.2) | 10 (3.1) | 50 (15.7) | 319 |
| *stl316/+ x stl336/+* | 335 (79.2) | 51 (12.1) | 37 (8.7) | 423 |

Therefore, distinct differences in the severity of phenotypes and rates of survival were observed between the more C-terminal located nonsense *stl316* allele and the more N-terminal *stl756* and *stl336* nonsense alleles. We reasoned that *stl756* and *stl336* were most likely null alleles, while *stl316* most likely represents a hypomorphic allele of *adamts9*. To test this, we performed pair-wise complementation testing of the candidate *adamt9*^*stl316*^ allele and the *adamts9*^*stl756*^ and *adamts9*^*stl336*^ alleles. We found that either the *adamts9*^*stl756*^ or the *adamts9*^*stl336*^ allele failed to complement the post-embryonic onset of caudal scoliosis commonly observed in the *adamts9*^*stl316*^ mutants (Table 3). However, in contrast to homozygous *adamts9*^*stl316/stl316*^ mutants, we observed consistently smaller sized fish with a higher incidence of misshapen eyes and more severe spine curvature phenotypes in both *adamts9* ^*stl316/stl756*^ and *adamts9* ^*stl316/stl336*^ transheterozygous mutant zebrafish (20 dpf) (Fig. 6 E, F; Table 3). We also observed a striking reduction in the mineralization of the spinal column in both *adamts9* ^*stl316/stl756*^ and *adamts9* ^*stl316/stl336*^ transheterozygous animals (Fig. 6 D, F). Such phenotypes were not observed in WT siblings (Fig. 6 B) or in *adamts9* ^*stl316/stl316*^ homozygous mutant fish (Fig. 5 E).

To address which tissues could be compromised in our *adamts9* mutant zebrafish, we utilized whole-mount *in situ* hybridization to determine the spatio-temporal patterns of *adamts9* gene expression. Human *ADAMTS9* is alternatively spliced toward the 3’ region. The RefSeq Adamts9 protein sequence currently available at Ensemble (GCA_000002035.3) lacks the terminal GON1 domain, which is annotated in human, *C. elegans*, and mouse Adamts9 protein (Kelwick et al., 2015). To determine if the WT zebrafish Adamts9 lacks the terminal GON1 domain and to generate a full-length *adamts9* transcript for *in situ* hybridization, we used 3’-Rapid Amplification of cDNA Ends (3’RACE) to ensure accurate cloning of 3’ end of the *adamts9* transcript. Sanger sequencing showed that zebrafish Adamts9 protein shares 48.7% identity with human ADAMTS9, however as predicted by the RefSeq database, our full length *adamts9* clone also lacked any sequence predicted to encode a C-terminal GON1 domain.

**Figure 6:**
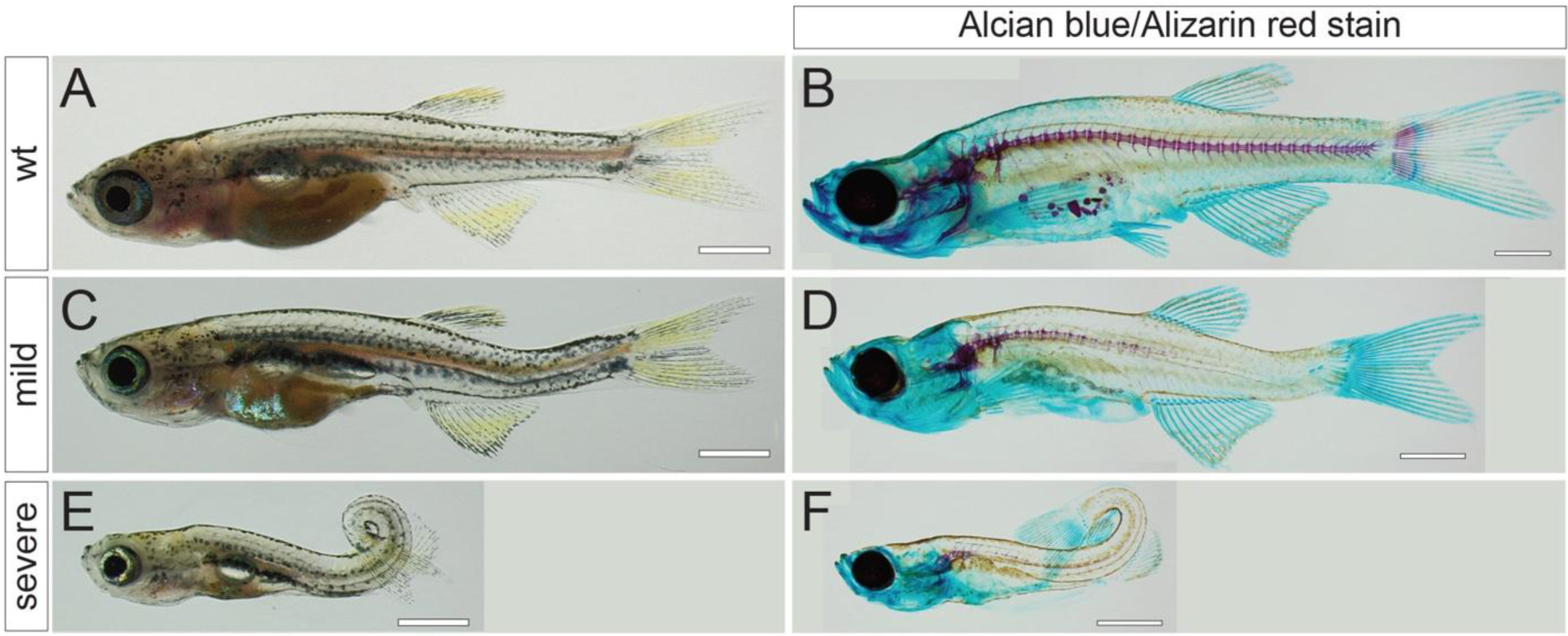
Transheterozygous *adamts9*^*stl756/stl316*^ mutants display severe body curvature phenotypes. **(A, C, E)** Bright field and **(B, D, F)** Alizarin-Red and Alcian-Blue stained skeletal preparations illustrate representative phenotypes from an *adamts9*^*stl756/+*^ x *adamts9*^*stl316/+*^ intercrosses. **(A, B)** wild type, **(C, D)** mild, and **(E, F)** severe phenotypes.

By *in situ* hybridization using full-length *adamts9* riboprobe we detected *adamts9* expression in the brain (red arrows, Supp. Fig. 7), the ciliary marginal zone of the eye, the lateral line structures, and a subset of cells spaced along the dorsal-ventral tips of each somite (yellow arrows, Supp. Fig. 7). We did not observe *adamts9* expression in the notochord or spine tissues, nor anywhere specific to the caudal portion of the animal at any stage we tested. Nor did we find any evidence for defects in the musculature of the posterior flank by transverse sections, stained with H&E or when assayed by immunohistochemistry using antibodies against Myosin Heavy Chain muscle markers of both Pan and Fast muscle markers (Supp. Fig. 8, 9). However, we did observe a consistent dilation of the central canal in transverse H&E sections of *adamts9*^*stl316/stl316*^ mutants compared to WT siblings (Supp. Fig. 9).

Given the dilated central canal in the *adamts9*^*stl316/stl316*^ zebrafish mutants, we next wanted to assess whether central canal structures were defective. Recent studies demonstrate that disruption of a central canal resident structure called the Reissner fiber (RF) results in defects of spine morphogenesis (Troutwine et al., 2020). To determine whether the RF is defective in *adamts9*^*stl316/stl316*^ mutants, we utilized the AFRU antiserum (Rodriguez et al., 1984) to stain zebrafish RF. We found no defects in RF morphology at 15 dpf (Figure 7), at the stage when caudal curvatures were obvious in *adamts9*^*stl316/stl316*^ mutants (Figure 5A). Altogether our genetic analysis highlights a critical role for Adamts9 for viability, eye development, and spine morphogenesis. Our analysis of a C-terminal truncation allele of Adamts9 implicates a novel mechanism controlling spine morphogenesis that is not due to disruption of the Reissner fiber.

**Figure 7:**
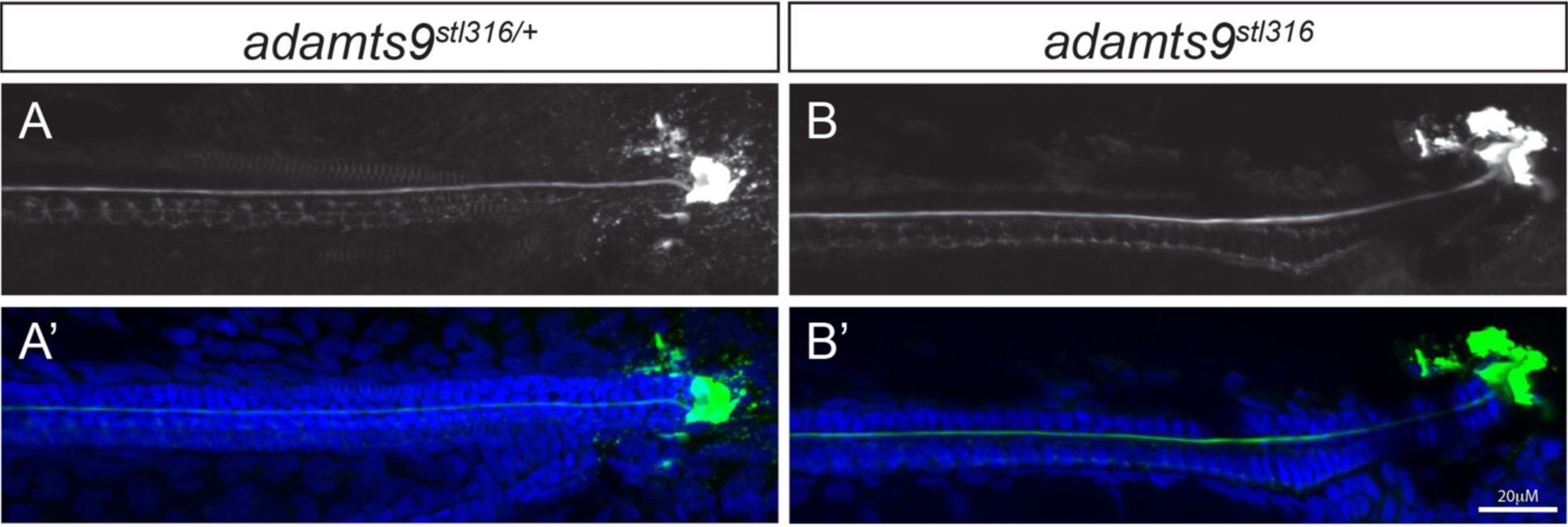
The Reissner fiber is not obviously affected in *adamts9*^*stl316/stl316*^ mutants. Confocal images of AFRU stained Reissner fiber (A, B) and merged images with DAPI-stained nuclei (A’, B’) in *adamts9*^*stl316/+*^ heterozygous (A, A’) and *adamts9*^*stl316/stl316*^ mutants (B, B’) at 15 dpf, which shows no obvious defects in Reissner fiber assembly.

## Discussion

Our forward genetic screen identified three dominant and 36 recessive adult viable mutations affecting spine development. We characterized four distinct phenotypic groups of adult-viable scoliosis mutants that model a variety of spine disorders observed in humans. Utilizing WGS, exome capture, and SSLP-based mapping approaches to map a subset of the mutant loci, we identified and confirmed several new candidate gene mutations affecting the spine morphogenesis in zebrafish.

The majority of mutants isolated, collectively called Group I, most closely resembled AIS in humans, in that the vertebral bodies are not malformed and are well segmented and the spine curvatures are found along the length of the spine with areas of spine rotation (Cheng et al., 2015). In contrast, group II mutants appeared generally stunted in body length. Some group II mutants displayed vertebral malformations and fusions, which resembles the pathology found in human congenital scoliosis patients (Giampietro, 2012). These vertebral malformations followed stereotyped embryonic defects in the notochord which resemble mutations affecting copper metabolism (Madsen and Gitlin, 2008) and collagen components of the extracellular notochord sheath (Christiansen et al., 2009; Gray et al., 2014). We isolated a single group III mutant that displayed a severe lordosis in the thoracic portion of the notochord at a time point when the mineralization of the spine is first occurring. Spontaneous thoracic lordosis is observed in sheep with an autosomal recessive inherited form of rickets (Dittmer et al., 2009). To our knowledge congenital thoracic lordosis is extremely rare in humans but has been observed (Winter et al., 1978). We identified several group IV mutants, which displayed a caudal localized lordosis. Similar dorsal flexion phenotypes are observed in *polycystin 2* mutant zebrafish and caused by increased collagen type-II protein synthesis and secretion around the notochord (Le Corre et al., 2014). Caudal lordosis is also associated with alterations in urotensin neuropeptide signaling from the cerebrospinal contacting neurons in the central canal and response of slow-twitch muscle fibers in the dorsal somites (Lu et al., 2020; Zhang et al., 2018). Mild lumbar lordosis is typical of the human spine morphology. Exaggerated lumbar hyperlordosis is an acquired disorder, associated with a diverse number of syndromes and disorders in human (https://www.ncbi.nlm.nih.gov/medgen/263149). Finally, we identified three dominant mutations which caused congenital scoliosis phenotypes such as shortened body stature and vertebral fusions, which is a common phenotype after mutation of type I collagen in human and in zebrafish (Gistelinck et al., 2018l; Henke et al., 2017).

Our pilot genetic screen revealed 39 recessive and dominant mutations effecting spine in just over 300 haploid mutagenized genomes, nearly 1 scoliosis mutation in 10 mutagenized genomes, a surprisingly high frequency. Our complementation testing with only a subset of recovered mutations showed at least 9 linkage groups from 16 tested mutations, suggesting the screen is still discovering new genes rather than alleles of previously found mutations. These observations indicate that numerous zebrafish genes, likely important for the regulation of multifactorial inputs are necessary for spine morphogenesis and maintenance of spine stability.

More in depth genetic analysis was largely focused on the group I mutants, where we identified a novel non-synonymous mutation of *kif6*, which was previously implicated in larval-onset scoliosis in zebrafish (Buchan et al., 2014; Konjikusic et al., 2018). However, this report of a new *kif6*^*stl95*^ allele highlights the importance of an invariant Leucine residue (*Danio rerio*: L243Q) located in the well-conserved switch II domain (K[L]HLVDLAGSERV) in the motor domain. At the same time, analysis of a human disease-related *KIF6* p.P293T variant (from the Baylor data) provides more evidence of the critical role of Kif6 function for spine morphogenesis in zebrafish. Altogether this genetic analysis of *kif6* in zebrafish underscores the functional importance of evolutionarily conserved amino acid residues in kinesins and strongly suggest that any non-synonymous mutation of these residues in Kinesins should be scrutinized as strong candidate causative mutations of undiagnosed human diseases, akin to evidence of *KIF1A* mutations underlying human neurological disorders (Chiba et al., 2019; Pennings et al., 2020).

We recently demonstrated that *kif6* mutants have a defect in Reissner Fiber (RF) formation after 1 dpf (Troutwine et al., 2020), yet how Kif6 controls RF assembly is still under investigation. Kif6 is critical in zebrafish and mouse for the formation of ependymal cell cilia (Konjikusic et al., 2018), which are thought to contribute to ‘near-wall’ laminar flow of the cerebrospinal fluid in the ventricles and spinal cord (Spassky and Meunier, 2017). Perhaps the dynamic regulation of laminar flow is critical for assembly of the RF. An alternative model might involve the known requirement for cilia dependent regulation of cerebrospinal fluid volume in mice (Banizs et al., 2005), which may affect the distribution of the RF material from the subcomissural organ in the brain. SCO-spondin is the major protein component of the RF in zebrafish (Cantaut-Belarif et al., 2018). More comprehensive analysis of the non-complementing Group IA mutants, *stl297* and *stl300*, found in this screen, demonstrated these are non-synonymous mutations of the *scospondin* gene, causing disassembly of the RF during larval development leading to AIS-like spine defects (Troutwine et al., 2020). Given the similarity of Group IA phenotypes and the molecular implications for these mutations causing instability of the RF during larval development, it will be intriguing to determine whether the onset of whole-body scoliosis in other Group I mutants is also associated with defects in ependymal cilia and/or RF formation/maintenance. We anticipate that comprehensive analysis of the RF in our collection of Group I mutants may reveal mutations which do not affect the RF, thus implicating a downstream or parallel pathway for the molecular control of spine morphogenesis for this particular pathology. Despite our strong indication for the importance of the RF for spine morphogenesis, whether the RF is present in humans or apes remains controversial. Regardless it is important to understand if the pathways involved in downstream mechanisms, i.e., involving sensorimotor feedback control of posture, are conserved between fish and humans. Recent work indicates a postural control mechanism is downstream and responsive to stimulation of CSF-contacting neurons by RF in zebrafish (Knafo and Wyart, 2018; Orts-Del’Immagine et al., 2020; Zhang et al., 2018). CSF-contacting neurons are conserved in vertebrates, suggesting that while RF may be absent in humans, these neurons still function in postural control, while their stimulation may be different. Further genetic analysis of the genes involved in this physiologic circuit will be useful in the analysis of scoliosis in humans.

We molecularly characterized the Group IV *stl316* allele as a nonsense mutation in the *adamts9* gene predicted to truncate the C-terminal portion of the encoded protein. Adamts9 is a multi-functional metalloprotease, which has several established roles, including in extracellular matrix (ECM) turnover of proteoglycans, such as aggrecan and versican (Dubail and Apte, 2015; Nandadasa et al., 2015); ciliogenesis (Choi et al., 2019; Nandadasa et al., 2019); and a protease-independent function in promoting endoplasmic reticulum to Golgi transport for a variety of secretory cargos (Yoshina et al., 2012). In mouse, mutation of *Adamts9* results in lethality prior to gastrulation, hampering analysis during post-natal development (Dubail et al., 2014).

However, recent analysis of *Adamts9* conditional loss of function has begun to demonstrate a variety of roles for this metalloprotease including umbilical cord and eye development (Dubail et al., 2016; Nandadasa et al., 2015), and implication of its role in nephronophthisis, coloboma, and ciliogenesis (Choi et al., 2019; Nandadasa et al., 2019).

Our screen identified two truncating mutations of the zebrafish *adamts9* gene, with different phenotypic spectra. The *adamts9*^*stl316*^ allele, displays caudal-localized body curvature during larval development and caudal scoliosis in adult zebrafish. In addition, we observed eye defects with low penetrance and variable expressivity. It will be interesting to compare the eye defects in zebrafish *adamts9*^*stl316/stl316*^ mutants with those observed in *Adamts9* haploinsufficent mice, which is implicated in Peters Plus syndrome in humans (Dubail et al., 2016). In contrast, the *adamts9*^*stl756*^ allele is an early-truncating nonsense mutation, which is mostly embryonic lethal. However, we did observe a few escapers, which displayed severe defects in jaw and eye development, scoliosis. As *adamts9* ^*stl756/stl316*^ transheterozygotes are viable adults manifesting scoliosis, we conclude that *adamts9*^*/stl316*^ is a hypomorphic allele. Our genetic analysis demonstrates that the C-terminal domain, containing Thrombospondin repeats (TSRs), is required for the normal morphogenesis of the spine, barbels, and eye but not for viability. For these reasons it is interesting to speculate that the TSRs domains of Adamts9 provide a critical signaling function for larval spine morphogenesis in zebrafish. The TSRs are a conserved motif found as one or more copies in numerous proteins including: Thrombospondins, Semaphorins, ADAMTS family proteins, Complement proteins, and Spondin proteins. TSR domains have been shown to bind extracellular matrix components such as decorin, collagen V and fibronectin, bind and activate TGF-β, and bind TSR synthetic peptides and are sufficient to stimulate neurite outgrowth (Adams and Tucker, 2000), which could alter peripheral innervations critical for spine proprioception. Alternately, the loss of TSR domains might alter interactions between ECM components and possibly be part of a cartilage/ECM development affecting the structural integrity of the spine. Instead, *adamts9* ^*stl756/stl316*^ and *adamts9* ^*stl316/stl316*^ may function as hypomorphs if the ribosome reads through the nonsense mutation to produce full-length protein. A final possibility is that, genetic compensation may be triggered by *stl316* but not by *stl756* allele (El-Brolosy et al., 2019; Rossi et al., 2015), and these alternative mechanisms warrant further analysis.

Interestingly, we found no evidence of any defects in the formation of the Reissner fiber in the hypomorphic *adamts9*^*stl316/stl316*^ mutant fish, which suggest that Adamts9 acts in a downstream pathway or in a parallel pathway to control spine morphogenesis. Recent studies in human and model systems highlight the importance of the extracellular matrix in the formation of the spine and the pathology of scoliosis (Wise et al., 2020).

It is also noteworthy that for the two of the three molecularly identified mutant loci: *kif6, scospondin* and *adamts9*, adult viable scoliosis phenotypes are caused by hypomorphic mutations in otherwise essential genes. We wonder if adult viability in a scoliosis phenotype in zebrafish requires mutations be partial functioning to permit life and whether this is relevant to human scoliosis. In addition to the importance of extracellular matrix, our studies indicate the importance of the nervous system and the Reissner fiber in scoliosis as well as, emphasize the RF independent regulation of spine morphogenesis.

These novel mutant zebrafish lines and their molecular characterization will provide a resource for new biological insights into spine morphogenesis in zebrafish. The pilot screen of 314 mutagenized haploid genomes yielded 39 loci affecting adult viable body straightness. Our studies suggest that a large number of genes are involved during spine morphogenesis in zebrafish. Complementation and mapping experiments indicate most of the mutations affect distinct genes, indicating the screen is far from saturation and continued screening will identify additional genes underlying normal spine development and scoliosis.

## Materials and Methods

### Fish Strains

The SAT WT strain was used in mutagenesis. Briefly, this strain originated as a AB:Tübingen hybrid strain. Each of the parental lines was extensively incrossed to maximize homozygosity at each gene locus (Kettleborough et al., 2013). All zebrafish studies and procedures were approved by the Animal Studies Committees at the University of Texas at Austin and the Institutional Animal Care and Use Committee at Washington University School of Medicine in St. Louis.

### Design of the Genetic Screen and Mutagenesis

Our screen was designed to recover zygotic dominant and recessive mutations in genes involved in adult viable spine disorders, namely congenital-type scoliosis including malformations of the vertebral units and spine curvatures; and larval onset scoliosis, displaying no discernable malformations of vertebral units at the onset of scoliosis, pathologically similar to adolescent idiopathic scoliosis in humans. Moreover, we wanted to provide a genetic resource to compare to targeted sequencing of human spine disorders. Male zebrafish were mutagenized with *N*-ethyl *N*-nitrosourea to induce mutations at an average specific locus rate of approximately one in 800 mutagenized genomes. A total of 1,229 F1 fish were screened for dominant adult viable phenotypes. Our adult F3 screen for recessive scoliosis phenotypes screened 314 F3 haploid genomes. The number of haploid mutagenized genomes screened was calculated according to Driever 1996 (Driever et al., 1996):

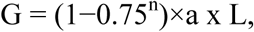

where G=haploid genomes analyzed. This assumes the F1 is heterozygous, the F2 is result of F1 crossed to WT, so that the F2 stock is composed of 1:1 of WT:heterozygous fish. The 0.75 term is the chance of an uninteresting WT/WT (0.25) or WT/het (0.50) pair in the cross. The term (1-0.75^n^) is the chance that the F2 cross is a pair of heterozygous parents. n= average number of crosses with more than 20 embryos per line. See that as the number of crosses increases, the chance of having a useful heterozygous pair in the cross approaches 1. The term a= number of mutagenized haploid genomes per family: a=1 for F1×AB; L= number of lines examined. Our numbers were 671 lines, an average of 3-4 clutches per family and F1xAB lines.

### Fish Husbandry and Mutagenesis

The zebrafish rearing and breeding protocol have been described by Solnica-Krezel (Solnica-Krezel et al., 1994). Adult male fish were mutagenized in 3 mM or 3.5 mM ENU in the presence of 10 mg/L MS-222 (tricaine) at 20-21°C for 4 or 6 times for 1-hour periods at weekly intervals. After 1 hour of mutagenesis mutagenized fish were carefully and quietly transferred to a tank with 10 mg/L MS-222 at17-19°C. After 6 hours fish were gently moved to fish water without MS-222. Two weeks after the end of ENU treatment, males were crossed to females at weekly intervals to build F2 families. On average 3-5 F3 clutches were reared per F2 family giving between ∼58-76% chance of observing a novel recessive mutation per family. In general, 60 fish were placed in a 1.1 Liter tank with a 500-micron screen baffle (Tecniplast) at 4-5 dpf. All tanks were hand-fed rotifers, for 8-12 days. After 13 days, tanks were placed on an automated feeding system (Tecniplast) of dry food 4 times a day at 20 mg per feeding, and 10 mL of artemia/rotifer mixture twice a day. Artemia cultures were started with an average egg concentration of 0.253 g/mL and average rotifer concentration was 1150 rotifers per mL. Automated feeding supported increased survival of homozygous mutants and increased density, and growth rate of our larval fish.

### Screening Procedure

All F2 families were screened for dominant adult viable phenotypes prior to incrossing, which was used to generate F3 families. F3 families were grown at a density of 50 larvae/ 1.1 L tank. All clutches were scored at 15 dpf and at 25-30 dpf to identify post-embryonic defects, particularly scoliosis, that effected 10-25% of the population. We screened progeny in tanks by eye and, in some cases, using a dissecting microscope. We confirmed all dominant mutations by outcrossing a single representative of the putative mutant to the original isotype strain (SAT). Multiple sperm samples from mutant fish as well as the F0 gametes were cryopreserved.

### Outcross and Complementation Testing of Recessive Mutations

A phenotypically representative, putatively F3 recessive mutant was outcrossed to usually, a WT SAT strain or rarely to an SJD mapping strain. In the rescreen, two or three F4 (presumably heterozygous mutant carriers) were incrossed and clutches of 40-80 individuals were grown to and screened at 25-30 dpf. The F4 parents were saved in a 1.1 L tank so that sperm from confirmed heterozygous carriers was cryopreserved according to (Matthews et al., 2018). To test for genetic complementation between different alleles, we performed crosses between mutants with similar phenotypes.

### Mapping using Whole Genome Sequencing

To map a mutation, a single F4 homozygous mutant was outcrossed to SJD or WIK to generate a mapping cross family. Several pairs of these animals, presumably heterozygous for the mutation, were intercrossed and F5 progeny were screened for heritable spine defects. DNA was extracted from 20 phenotypically WT and 20 mutant larvae at 30 dpf. Equimolar concentration of DNA was taken from each individual and pooled into mutant and WT samples and sent to the Genome Technology Access Center (GTAC, Washington School of Medicine) for whole genome sequencing. An in-house analysis pipeline was used to determine the region of homozygosity segregating with the mutant pool in contrast to the WT sibling pool (1 WT: 2 heterozygotes). A SNP subtraction analysis using other whole genome sequencing (WGS) datasets from this screen was used to narrow the number of candidate SNPs.

### Exome Capture Sequencing and DNA preparation

DNA for whole exome capture sequencing was extracted from 20 phenotypic animal or their WT siblings (adult or embryo, generated as described above) using the DNeasy kit (Qiagen) by the method described in (Sanchez et al., 2017). Automated dual indexed libraries were constructed with 250 ng of genomic DNA utilizing the KAPA HTP Library Kit (KAPA Biosystems) on the SciClone NGS instrument (Perkin Elmer) targeting 250bp inserts. Up to 15 libraries were pooled pre-capture generating a 5µg library pool. The library pool was hybridized with a custom Nimblegen probe set (Roche), targeting protein-coding regions and non-coding RNAs (lncRNAs and miRNAs), resulting in ∼42Mb of target space. The concentration of the captured library pool was accurately determined through qPCR according to the manufacturer’s protocol (KAPA Biosystems) to produce cluster counts appropriate for the Illumina NovaSeq6000 platform. 2×150 read pairs were generated per library pool yielding an average of 9Gb of data per sample. Approximately 50x mean target coverage was achieved.

### *adamts9* Cloning and Genotyping

cDNA was synthesized using the ProtoScriptII First Strand cDNA synthesis kit (NEB E6560S). Cloning of the full length *admats9* from pooled cDNA from 3, 5, and 15 dpf AB*zebrafish, due to issues with cloning a full-length cDNA of the *Danio rerio admats9* gene (ENSDART00000109440) we used a two PCR amplification, using LATaq (Takara), to obtain a full-length *adamts9* cDNA. The 1st amplification was done using a foward primer (5’-AATGCACTAGCTTCAACGGGG-3’) and reverse primer (5’-ATGAGCGTACCAAGCTCCA-3’) primer set. This amplicon was TA cloned (pCRII-TOPO; ThermoFisher) and Sanger sequenced. This vector containing a 5’ truncated *adamts9* cDNA with a small fragment of the 3’ UTR was used as a template for a 2nd PCR to add the truncated 5’-*adamts9* fragment and the 3’ stop site using a forward primer (5’-ATGGTTTTGTTCTCCTGGGGAATTAGTTTTTTAGTATTACTCTCTGATCTAATGAATGCACTAGCTTCAACGGGGAGACTGCGAGGTTCAG-3’) and a reverse primer (5’-TTATGAGCGTACCAAGCTCCATTGGTCTGTGC-3’) and was TA cloned and sequenced. The amplicon was then TA-cloned into a dual promoter pCR TOPO-II vector (ThermoFisher). To ensure we had the full-length transcript we performed 3’RACE (SMARTer RACE Takara) using a forward primer 5’-GGAGCACAGACCAATGGAGC-3’.

The *stl316* mutation was sequenced using forward primer 5’-GAGAATCAACTGCACTGACCGC-3’ and reverse primer 5’-CTGTGTTGCATGACTCAGTGATCGC-3’. To genotype individual *adamts9*^*stl756*^ larvae for phenotypic analyses, we used the following primers to amplify a 174 bp fragment of adamts9 from genomic DNA: 5’-CTCGAGTCTGGATTTATTGC -3’ and 5’ GGAGCAGAGGCTGATTACT - 3’.

The ENU-induced mutation in *adamts9*^*stl756*^ introduced an MseI restriction enzyme recognition site that was used for genotyping. Following restriction digest using MseI, which generates a 27 bp product and a 147 bp product in mutant embryos but fails to cut WT, digested DNA was resolved on a 3% agarose gel. To genotype individual *adamts9*^*stl336*^ animals for phenotype analyses, we used the following primers to amplify a 252 bp fragment of the *adamts9* locus from genomic DNA: 5’-CTAGCTGAGCTG GGTACAGT -3’ and 5’-ACACTCAGGCCACATTTAGA -3’.

The TALEN induced mutation in *adamts9*^*stl336*^ disrupted an HphI restriction site, such that restriction enzyme digest of the aforementioned PCR product results in a 178 bp product and a 74 bp product in WT animals but fails to cut mutant. Digested DNA was resolved on a 3% agarose gel.

### *kif6 P293T* Genome Editing and Genotyping

The *kif6*^*dp20*^ mutant zebrafish were developed using CRISPR-Cas9-mediated genome editing. Using the CHOP-CHOP online tool (Labun et al., 2016), we identified a suitable 20-nucleotide site (GGTCATCATGGAATTGCGGTAGG) targeting exon 8 of Danio *kif6* (ENSDART00000103662.6) in order to generate non-synonymous p.P293T mutant allele. The gene specific and universal tracrRNA oligonucleotides (S1 Table) were annealed, filled in with CloneAmp HiFi PCR premix, column purified, and directly used for *in vitro* transcription of single-guide RNAs (sgRNAs) with a T7 Polymerase mix (M0255A NEB). All sgRNA reactions were treated with RNAse free-DNAse. We utilized a ssDNA oligonucleotide (S1 Table) to insert the desired base-pair changes, and a synonymous mutation which introduced an EcoRI site for genotyping (Supp. Fig. 2C). The *kif6* p.P293T donor and locus specific *kif6* sgRNA and purified Cas9 protein (Alt-R S.p. Cas9 Nuclease V3, IDT) were injected at the 1-cell stage. We confirmed segregation of the *kif6*^*dp20*^ allele using several methods: allele-specific PCR, or EcoRI (NEB) digestion of the *Kif6* exon14 amplicon, and Sanger sequencing of heterozygous carriers (Supp. Fig. 2C, D).

### AFRU immunofluorescence of Reissner Fiber

Embryos and larvae were fixed and processed as in (Troutwine et al., 2020).

## Acknowledgements

We would like to acknowledge inspiration from the The Simpsons - *The Quetzlzacatenango* (S8E9) for the original *pimento locura* allele name for *adamts9*^*stl316*^. Research reported in this publication was supported by the National Institute of Arthritis and Musculoskeletal and Skin Diseases of the National Institutes of Health under Award Number R01AR072009 (R.S.G.) and the National Institute of Child & Human Development 1P01HD084387 (L.S-K.). This publication was made possible in part by Grant Number UL1 RR024992 from the NIH-National Center for Research Resources (NCRR). Declarations of interest: none

