## supplemental figures for "Postembryonic screen for mutations affecting spine development in zebrafish"

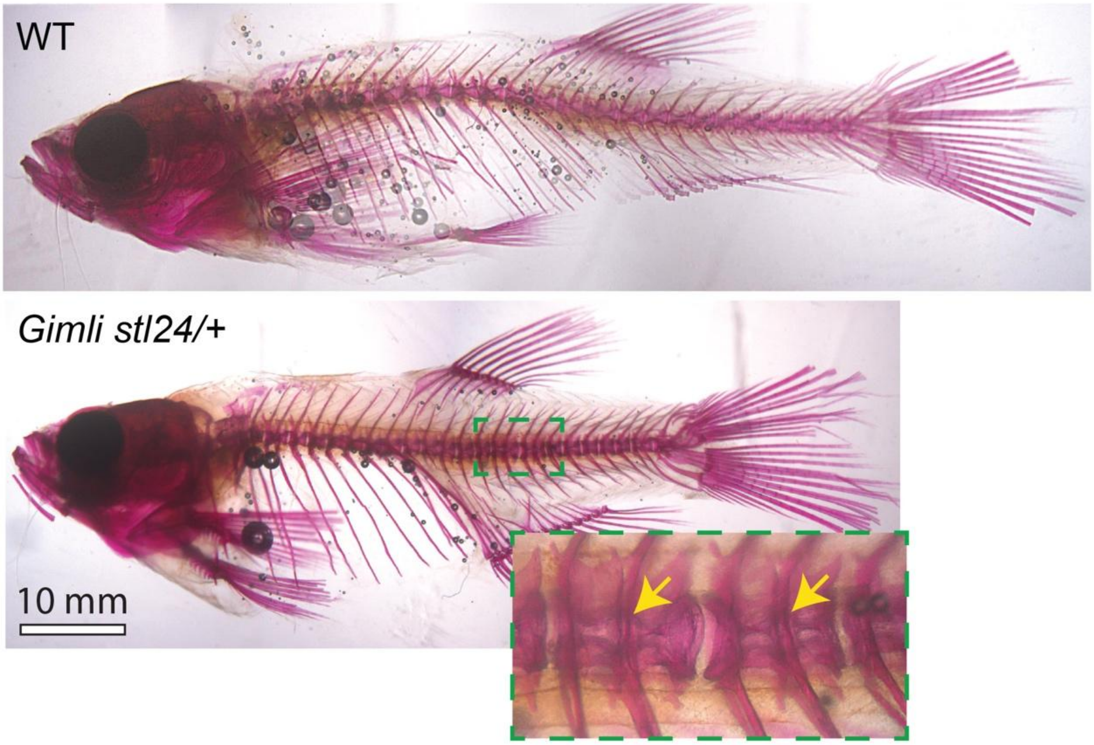


**Supplemental Figure 1: Representative group V dominant mutant displaying vertebral fusions.** Alizarin-red stained skeletal preparations.

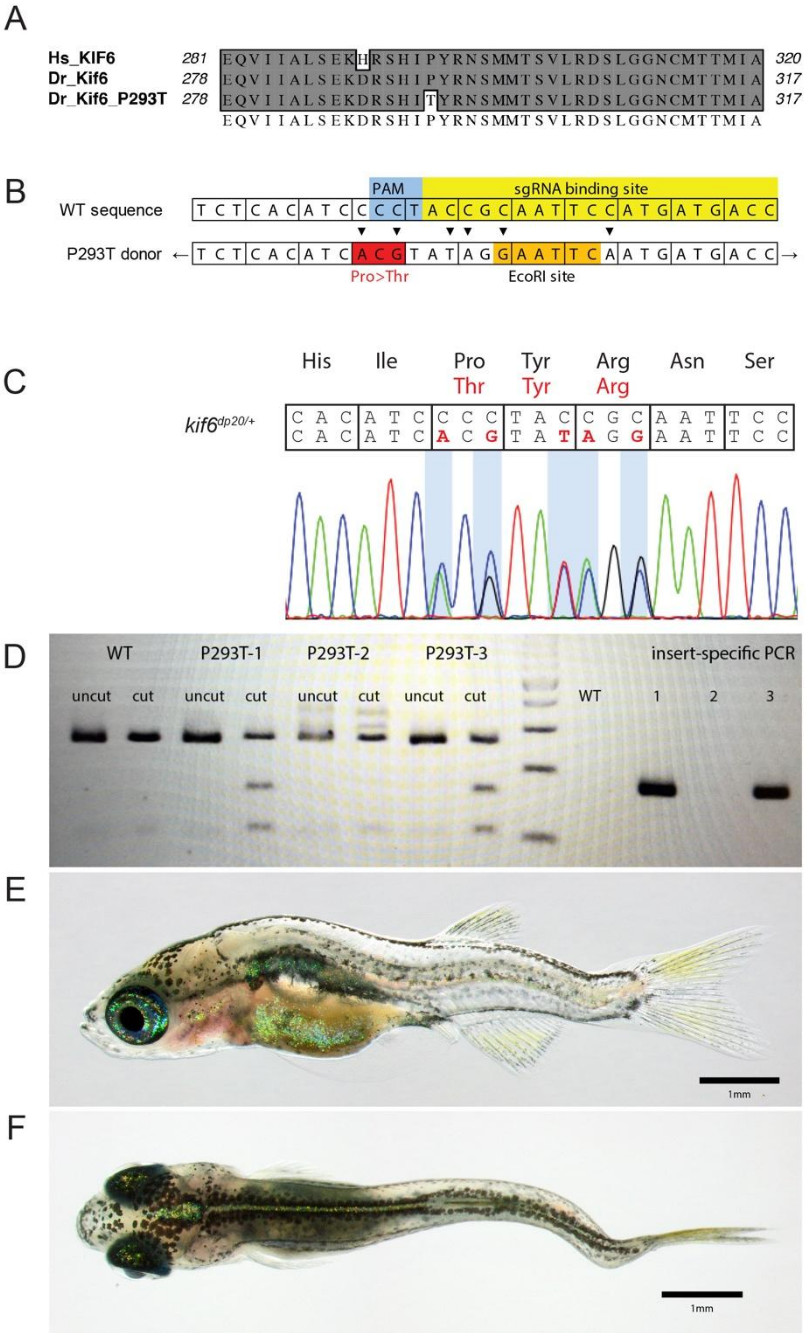


**Supplemental Figure 2: Generation of a novel *kif6* p.P293T mutant zebrafish. (A)** Clustal W alignment of Human and Danio Kif6 protein to highlight the P293T mutation **(B)** Schematic of changes for donor editing of the $kif6^{dp20}$ allele. **(C)** Sanger sequence of a stable heterozygous $kif6^{dp20}$ mutant zebrafish. **(D)** Representative EcoR1 and allele specific genotyping of the *kif6*$^{dp20}$ allele **(E, F)** Representative phenotypes of a homozygous *kif6*$^{dp20}$ mutant zebrafish at 30 dpf.

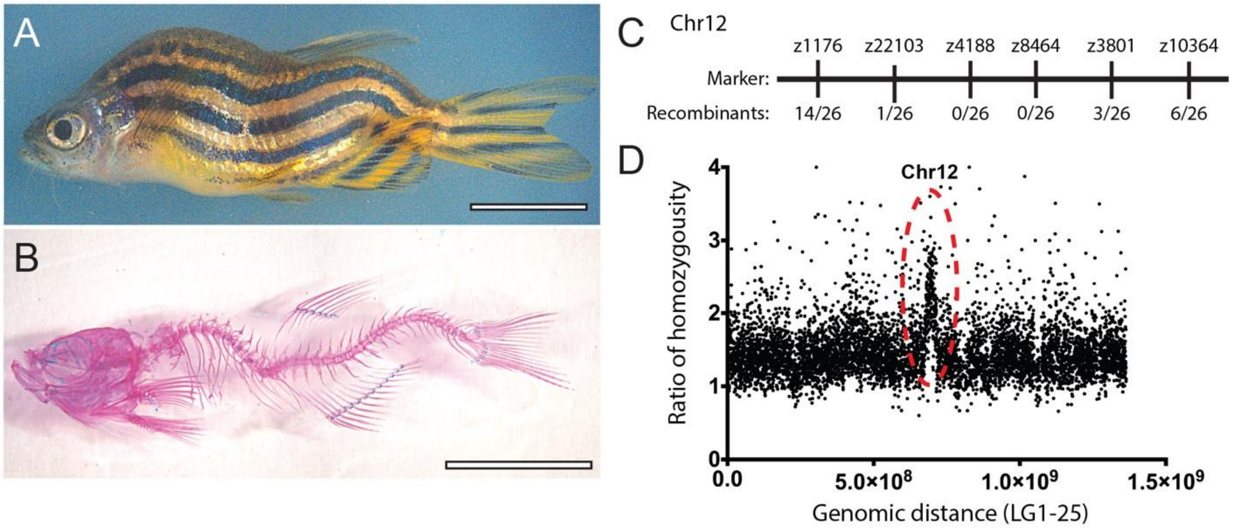


**Supplemental Figure 3: The *falkor* scoliosis mutant. (A)** Bright field and **(B)** Alizarin-Red/ Alcian-Blue stained skeletal preparations illustrate the homozygous *falkor* mutant phenotype. **(C)** Meiotic mapping places the mutation on Chr. 12 in the neighborhood of z4188 and z3801. **(D)** Mapping by regions of homozygosity suggests a causative mutation in *korken* gene on Chr. 12.

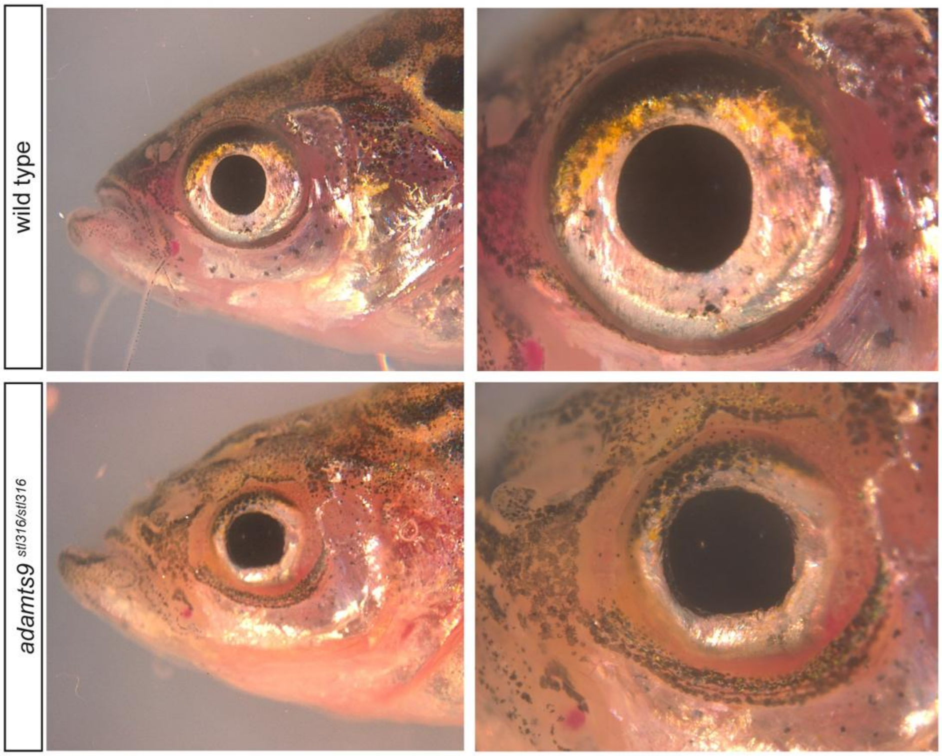


**Supplemental Figure 4:** ***adamts9*** $^{\mathbf{\textit{stl316/stl316}}}$ **mutant zebrafish display defects in eye development.**

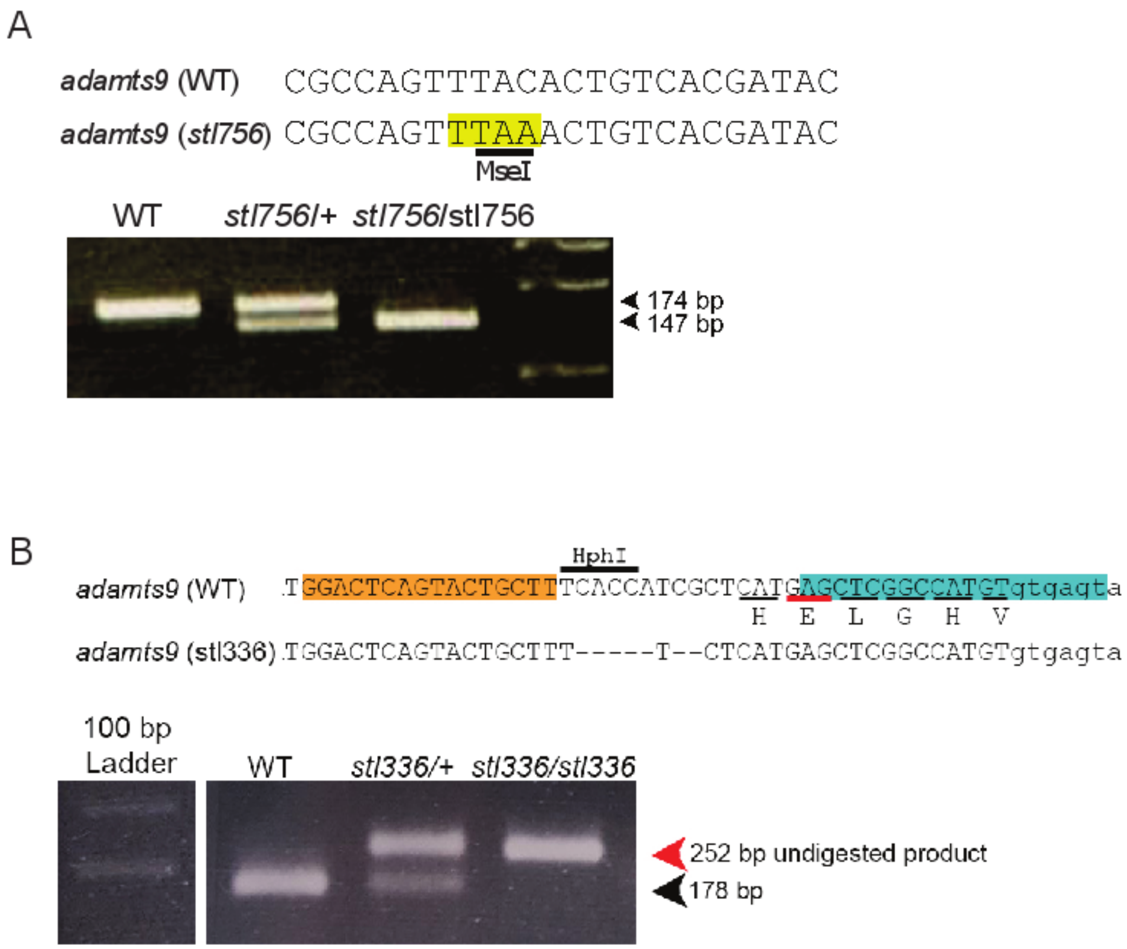


**Supplemental Figure 5: Genotyping methods for *adamts9*$^{stl756/stl756}$ and *adamts9*$^{stl336/stl336}$ mutants. (A)** Illustration of ENU-derived *adamts9*$^{stl756}$ mutation and representative genotyping by MseI digestion. **(B)** Illustration of the TALEN-derived *adamts9*$^{stl336}$ mutation and representative genotyping by HphI digestion.

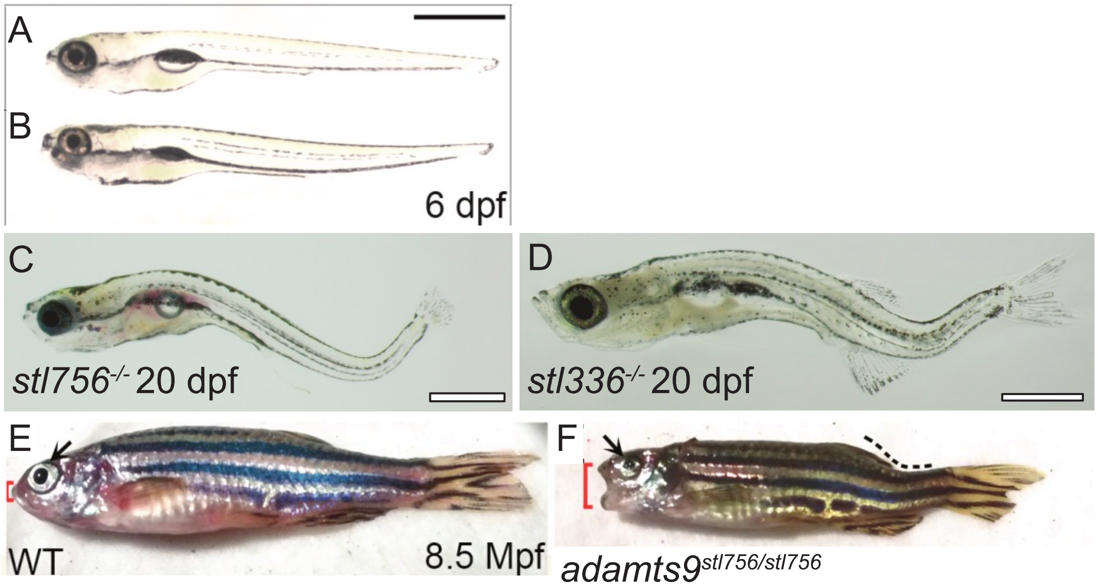


**Supplemental Figure 6: Severe phenotypes associated with the N-terminal truncation mutants of *adamts9.*** **(A-F)** Bright field imaging shows no phenotypes associated with *adamts9*$^{stl756}$ at 6 dpf. However, severe curvatures of the spine and eye defects are observed by 20 dpf for both **(C)** *adamts9*$^{stl756}$ and **(D)** *adamts9*$^{stl336.}$ **(F)** While most of these mutants are larval lethal, rare escapers can survive to adulthood but display severe defects in eye development (black arrow), spine curvature (dotted line), and defects of the jaw (red bracket), **(E)** not observed in WT siblings.

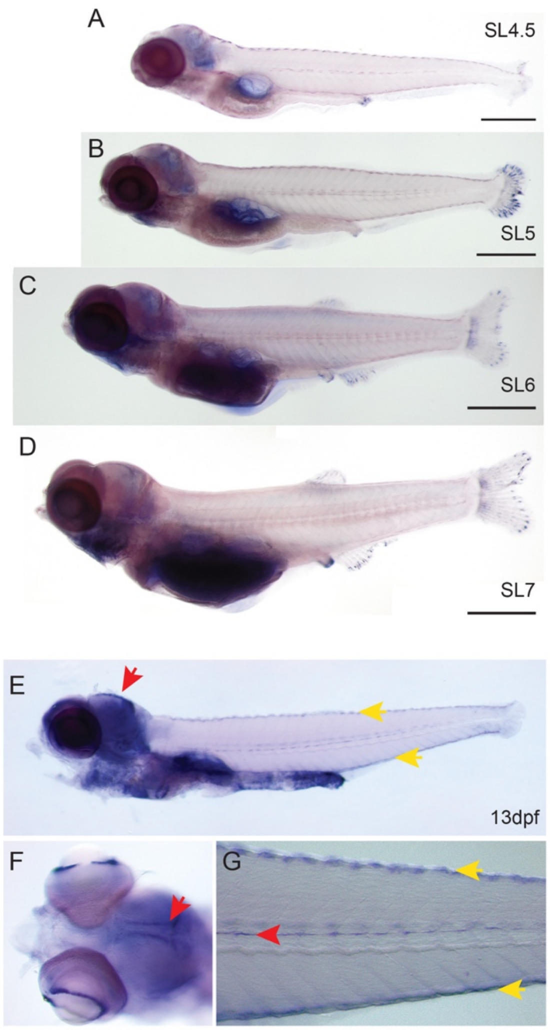


**Supplemental Figure 7: In situ hybridization of *adamts9* (A-D)** Lightly-stained staged series from WT fish showing *adamts9* expression in the head, cloaca, and at the tips of growing fins rays. **(E-G)** Extended staining at 13 dpf highlights specific *adamts9* expression in the head in bilateral strips in the brain (red arrows; **E, F**), staining of the melanocytes (yellow arrows), in the ciliated marginal zone of the eyes **(F)** and in the lateral line **(G)**.

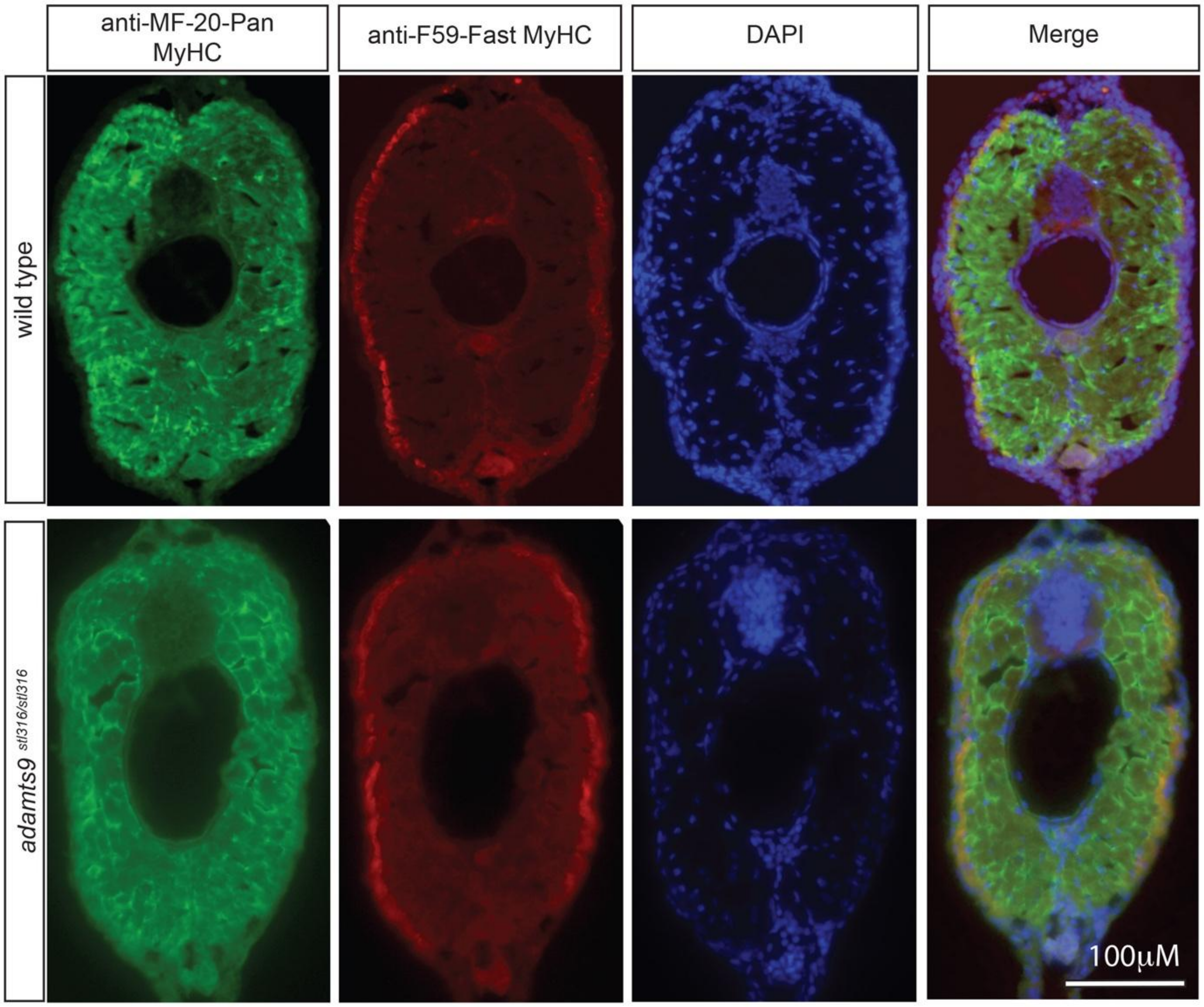


**Supplemental Figure 8: Muscle specific staining reveals no alterations in musculature in adult *adamts9*$^{stl316/stl316}$ mutants.**

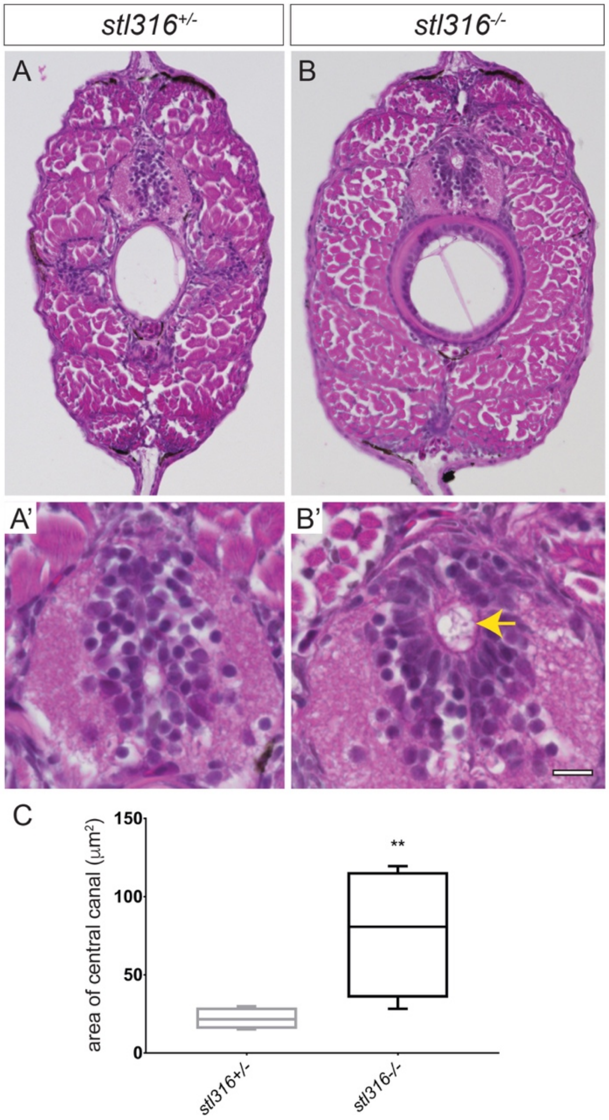


**Supplemental Figure 9: H&E staining of adult *adamts9*$^{stl316/stl316}$ mutants reveals a consistent expansion of the central canal (A-B')** Bright field of H&E stained transverse sections from heterozygous **(A, A')** and homozygous *adamts9*$^{stl316}$ mutants **(B, B')** at 90 dpf. **(C)** Graph of central canal area as box plot with SD and mean demonstrating a significant ($p>0.01$) increase in the central canal area in homozygous *adamts9*$^{stl316}$ mutants (yellow arrow) compared to heterozygous carriers.
